# ATP-dependent remodeling of chromatin condensates uncovers distinct mesoscale effects of two remodelers

**DOI:** 10.1101/2024.09.10.611504

**Authors:** Camille Moore, Emily Wong, Upneet Kaur, Un Seng Chio, Ziling Zhou, Megan Ostrowski, Ke Wu, Iryna Irkliyenko, Sean Wang, Vijay Ramani, Geeta J Narlikar

## Abstract

ATP-dependent chromatin remodeling enzymes mobilize nucleosomes, but how such mobilization affects chromatin condensation is unclear. Here, we investigate effects of two major remodelers, ACF and RSC using chromatin condensates and single-molecule footprinting. We find that both remodelers inhibit the formation of condensed chromatin. However, the remodelers have distinct effects on pre-formed chromatin condensates. ACF spaces nucleosomes without de-condensing the chromatin, explaining how ACF maintains nucleosome organization in transcriptionally repressed genomic regions. In contrast, RSC catalyzes ATP-dependent de-condensation of chromatin. Surprisingly, RSC also drives micron-scale movements of entire condensates. These newly uncovered activities of RSC explain its central role in transcriptional activation. The biological importance of remodelers may thus reflect both their effects on nucleosome mobilization and the corresponding consequences on chromatin dynamics at the mesoscale.

## Introduction

Within the nucleus, the level of chromatin condensation typically correlates with transcription repression, such that transcriptionally repressed heterochromatin is more condensed than transcriptionally active euchromatin (*1, 2*). ATP-dependent chromatin remodeling enzymes (*i*.*e*. ‘remodelers’) play critical roles in the formation and maintenance of these distinct chromatin domains(*3-5*). Remodelers drive changes in chromatin states by acting on the smallest unit of chromatin, a nucleosome, which consists of ∼ 140 bp of DNA wrapped around an octamer of histone proteins(*6*).

Substantial biochemical work has shown that remodelers can slide, disassemble, deform, and space nucleosomes, and that each remodeler has a unique impact on nucleosomes(*3, 4*). For example, the imitation switch (ISWI) class remodeler ACF has been shown to generate the evenly spaced nucleosome architecture found in heterochromatin(*7-11*). In comparison, the switch/sucrose non-fermentable (SWI/SNF) class remodeler RSC, which slides, deforms and disassembles nucleosomes, is critical for enabling DNA access in euchromatin(*12-15*). Most of our mechanistic understanding of remodeler action derives from studies at the nucleosome scale. At a genomic scale, ChIP based and genetic studies have shown correlations between the presence of specific remodelers and changes in chromatin organization in cells, and in vitro Micro-C studies have suggested that remodeler-driven nucleosome positions may promote formation of chromatin domains (*16-20*). Yet, how remodeler action at the nucleosome scale affects chromatin dynamics at a mesoscale remains an open question.

A simple prediction is that ATP-driven nucleosome mobilization disrupts inter-nucleosomal interactions, resulting in local chromatin de-condensation. Thus, remodelers may act as molecular “stir-bars” (*21*). Additionally, in cells, remodelers must operate within a crowded chromatin environment with estimated nucleosome concentrations of ∼100 uM or higher(*22*). How chromatin remodelers act in such a crowded environment is poorly understood. Recent studies have shown that chromatin condenses into phase-separated droplets *in vitro* that have comparable nucleosome concentrations to those within the nucleus(*23, 24*). Here, we build on these studies to ask how two key remodelers, ACF and RSC, which carry out substantially different transformations of a nucleosome, contend with a crowded chromatin environment. Further, as chromatin varies in nucleosome density and spacing in cells, we also investigate how nucleosome spacing and density affect chromatin condensation.

To investigate the interplay between condensed chromatin and remodelers, we combine single-molecule footprinting of chromatin fibers reconstituted on a native DNA sequence (single-molecule adenine methylated oligonucleosome sequencing assay of chromatin accessibility on assembled templates) (SAMOSA-ChAAT)(*25*) with confocal imaging of chromatin condensates. We find that irregular nucleosome spacing substantially increases the viscosity of chromatin condensates, but ACF and RSC can still act on nucleosomes within this environment. Contrary to simple predictions, ACF remodeling does not decondense chromatin, whereas RSC remodeling does. Additionally, unlike ACF, RSC activity promotes micron-scale motions of entire condensates. Our method for assaying both single-fiber remodeling and condensate properties enables insight into the mesoscale consequences of nucleosome remodeling. More generally, our findings demonstrate how remodeling activities that differ at the nucleosome scale can differentially change the chromatin environment at the mesoscale.

## Results

### Irregularly-spaced nucleosome arrays form viscous chromatin condensates

Previous studies have shown that chromatin assembled on DNA containing evenly spaced artificial 601 nucleosome positioning sequences forms phase-separated condensates under physiologically relevant buffer conditions(*23, 24, 26*). To investigate the phase-separation properties of chromatin assembled on a native sequence with weaker nucleosome positioning sequences, we assembled chromatin on a 3.5kb DNA sequence from the 5’ end of mouse gene *Cyp3a11* (sequence ‘S3’, in relation to sequences S1 and S2 studied previously (*25*)). Chromatin was assembled on fluorescently end labeled S3 and imaged by confocal microscopy under physiologically relevant buffer conditions (100mM KCl, 1.5mM free Mg^2+^)(Fig 1A)(*27*). Single-molecule nucleosome positions were determined as previously (*25, 28, 29*), by inferring methylase inaccessible footprints on individual molecules directly from PacBio sequencing data using a neural network-hidden Markov model. In contrast to templates assembled from on the evenly spaced 601 sequences (*28*) nucleosome positions assembled on S3 were irregular, consistent with observations for nucleosomes assembled on other physiological sequences(*25*).

**Figure 1.**
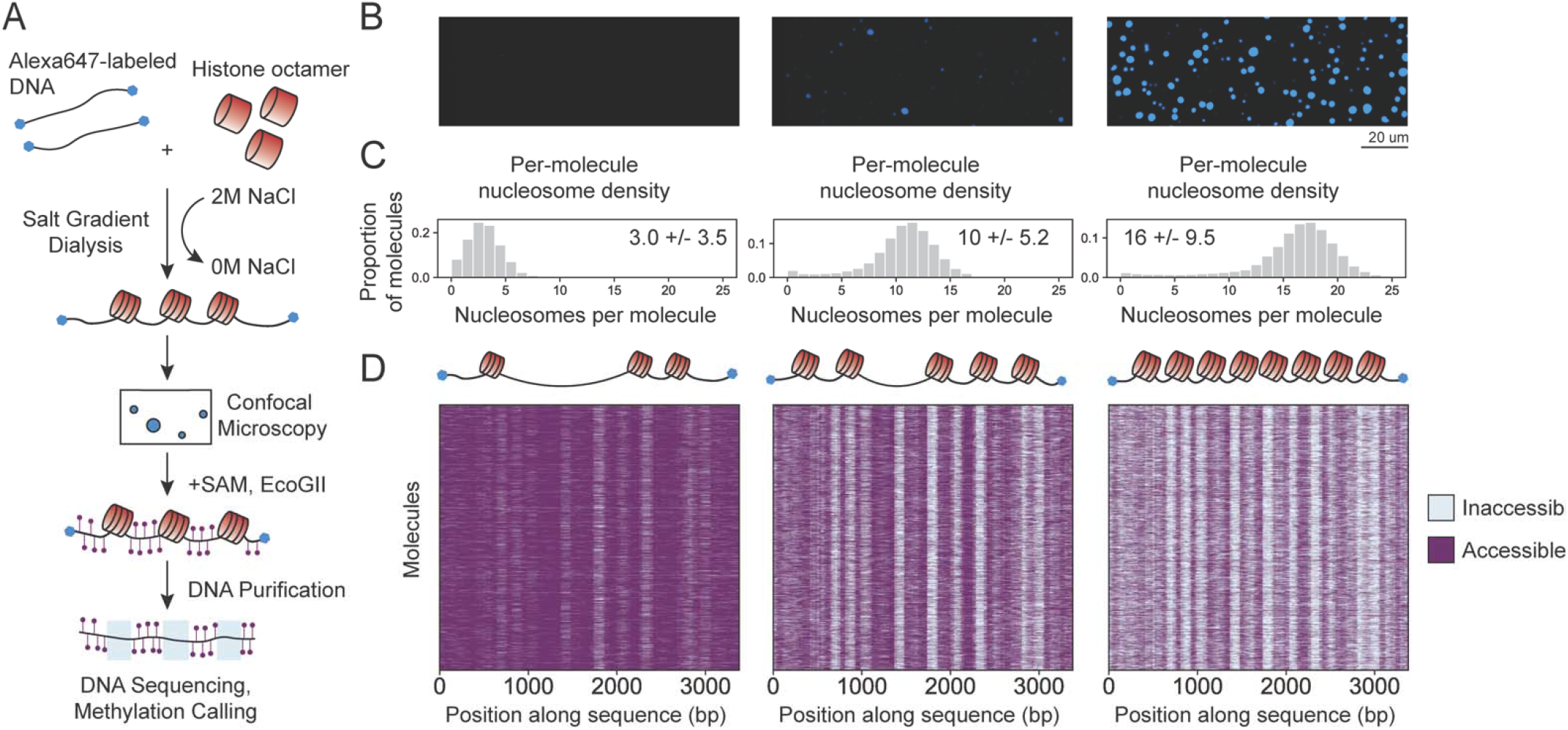
Nucleosome density drives chromatin condensate formation. (**A**) Scheme of experimental workflow. (**B**) Confocal images of chromatin reactions with varying histone octamer concentration showing Alexa647 fluorescence. (**C**) Per-molecule nucleosome density for each chromatin assembly shown in B. Inset is median nucleosomes per molecule and variance. (**D**) Heatmaps of accessible and inaccessible bases for 3000 molecules per condition. Cartoon chromatin molecules illustrate that each row in the heatmap represents a chromatin molecule and the number of nucleosomes per molecule increases according to octamer concentration.

Chromatin condensates are formed and stabilized by inter-nucleosomal interactions (*23*). We therefore hypothesized that increasing the number of nucleosomes per molecule (i.e. nucleosome density) on our irregularly spaced genomic sequence would promote chromatin condensation. Increasing the density of nucleosomes by varying the histone octamer:DNA ratio during assembly promoted phase separation, consistent with inter-nucleosomal interactions driving chromatin condensation (Fig 1B). In every assembly, we observed a distribution of nucleosomes per DNA template (Fig 1C and 1D), as previously observed(*25*). The median of this distribution shifted depending on how much histone octamer was used. Using the nucleosome density and the DNA concentration in the condensates, we calculated the concentration of nucleosomes in the medium and high-density chromatin condensates to be 7.6 +/-4 uM and 40 +/-23.5 uM, respectively (Fig S1A, see Methods). The nucleosome concentration in the high-density chromatin is comparable to concentrations measured by other groups (*23*), and that estimated within the nucleus (*22*).

We next investigated the dynamics of chromatin within the condensates by measuring the Fluorescence Recovery After Photobleaching (FRAP). Regularly spaced nucleosome arrays assembled on comparably long DNA templates containing 601 repeats (2 kb-3.3 kb) have been shown to display substantial FRAP within ten minutes, a feature we also recapitulated (Fig S2) (*26*). In contrast, we saw minimal fluorescence recovery within the S3 condensates ten minutes after photobleaching under all conditions, including a condition tested previously for the evenly spaced arrays (Fig S1B, Fig S2B)(*26*). As a complementary assay for chromatin dynamics, we performed a droplet mixing experiment using chromatin that was end-labeled with either AlexaFluor647 or AlexaFluor555 (Fig S1C). Upon mixing, we observed distinct, single-colored chromatin territories, indicating that these droplets were viscous, with low internal mixing (Fig S1C). However, the merged droplets were spherical, indicating that fused chromatin condensates do coalesce to minimize surface tension. Overall, our results suggest that irregular nucleosome spacing increases inter-fiber interactions inside chromatin condensates compared to regular spacing.

Given the high viscosity of the condensates, we wondered whether chromatin remodelers could access their nucleosomal substrates. We therefore investigated how the addition of remodeler affected (1) the formation of chromatin condensates and (2) the properties of pre-formed chromatin condensates. In both cases, we concomitantly assayed remodeling activity using SAMOSA-ChAAT.

### ACF inhibits formation of chromatin condensates in an ATP-independent manner

ACF generates regular chromatin arrays *in vivo* and *in vitro* and catalyzes nucleosome-spacing by sensing flanking DNA lengths(*25, 30*). To investigate how ACF regulates chromatin condensation, we mixed ACF with chromatin containing a median of 13 nucleosomes per template (Fig 2A). In the presence of ATP, ACF spaced nucleosomes in a density-dependent manner. The distance between nucleosomes was regular for a given DNA molecule, but varied from molecule to molecule depending on nucleosome density (Fig 2B, Fig S3). This observation is consistent with our previous work demonstrating density-dependent spacing by ACF on different genomic templates (*25*). Nucleosome footprints in reactions with ADP did not change relative to control chromatin (Fig 2D).

**Figure 2.**
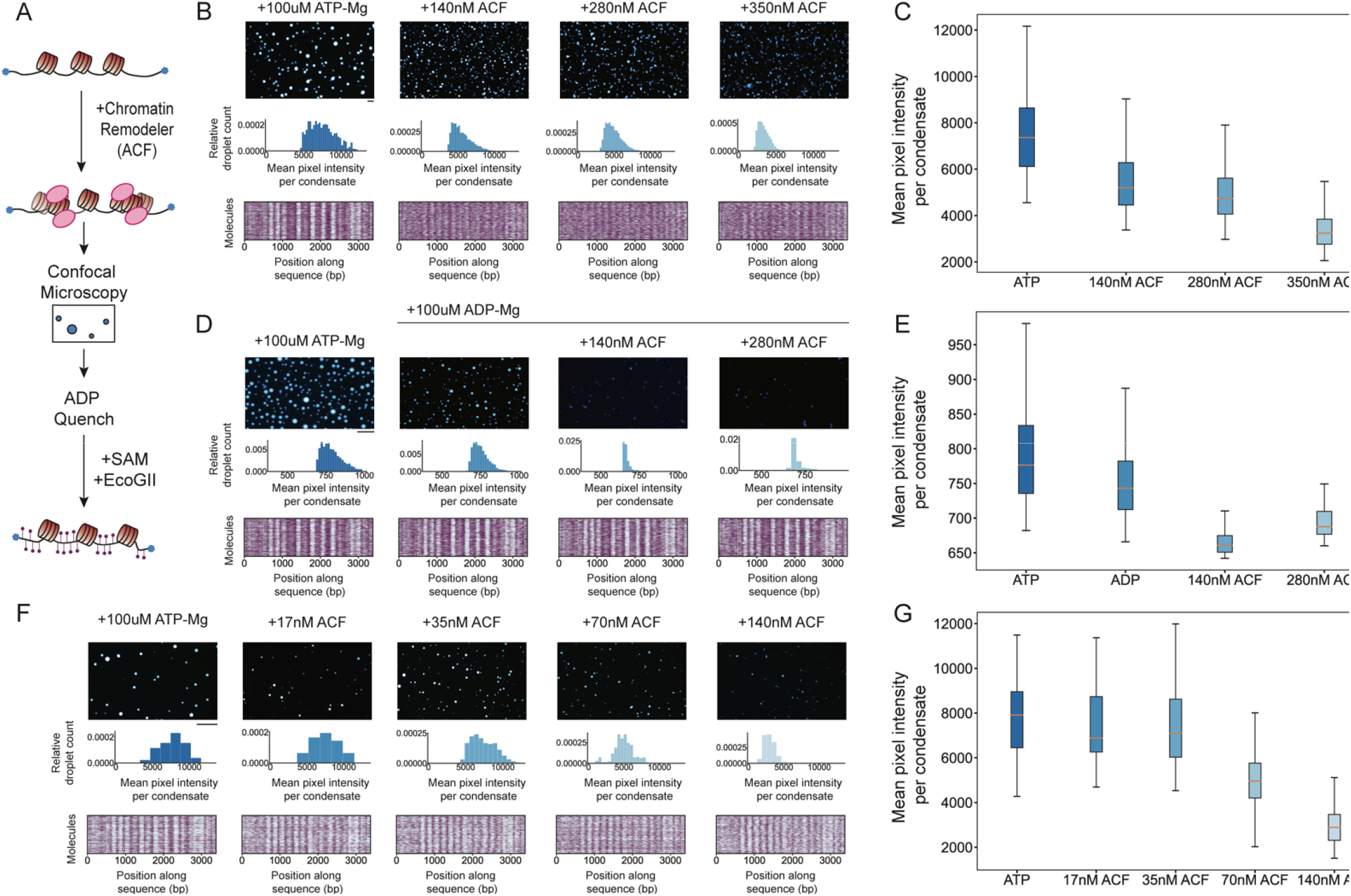
ACF prevents chromatin condensate formation in an ATP-independent manner. (**A**) Scheme of premix experimental workflow. (**B**) Titration of ACF in the presence of ATP-Mg. Confocal images showing Alexa647 fluorescence. Histograms of mean pixel intensity per condensate with histogram bin number equal to 0.1*number of condensates per condition. Heatmaps of accessible and inaccessible bases for 1000 molecules per condition. (**C**) Boxplot of mean pixel per droplet intensities from B. (**D**) Titration of ACF in the presence of ADP. Widefield images showing Alexa647 fluorescence, histograms and heatmaps as in B. Lower intensity threshold was increased for 140 and 280 nM samples for better visualization of condensates against background. (**E**) Boxplot of mean pixel intensities per condensate from D. (**F**) Titration of lower concentrations of ACF in the presence of ATP-Mg. Confocal images, histograms, and heatmaps as in B. (**G**) Boxplot of mean pixel intensities per condensate from F. All scale bars are 20 um.

ACF inhibited the formation of chromatin condensates, and condensates that did form were less intense and more irregularly shaped (Fig 2B-C). ACF had a similar effect in the presence of ADP or ATP, suggesting that binding by ACF to the arrays, rather than its nucleosome spacing activity, was responsible for inhibiting condensation (Fig 2B-E). Consistent with a binding effect, ACF inhibited chromatin condensation in a concentration dependent manner, having the largest inhibitory effect at stochiometric ratios of ACF:nucleosome (Fig 2F-G). These results rule out the model that ATP-driven nucleosome mobilization by ACF prevents chromatin condensation. Rather, the data suggest that ACF binding occludes the nucleosomal surfaces that participate in inter-nucleosomal interactions.

### ACF remodels nucleosomes in pre-formed condensates without dissolving them

Given our observation that stoichiometric amounts of ACF inhibit formation of chromatin condensates, we wondered if ACF would also dissolve pre-formed chromatin condensates. To test this possibility, we added ACF into a mixture containing preformed chromatin condensates (Fig 3A). Chromatin condensates persisted after adding ACF, regardless of whether we added 350 nM or 800 nM ACF to condensates generated from chromatin with ∼300 nM nucleosomes (Fig 3B, Fig S3C). These findings contrast with those in Fig. 2 showing that pre-mixing 350 nM ACF with chromatin inhibits condensate formation. A simple explanation of this result is that only a minimal amount of ACF entered the pre-formed condensates.

**Figure 3.**
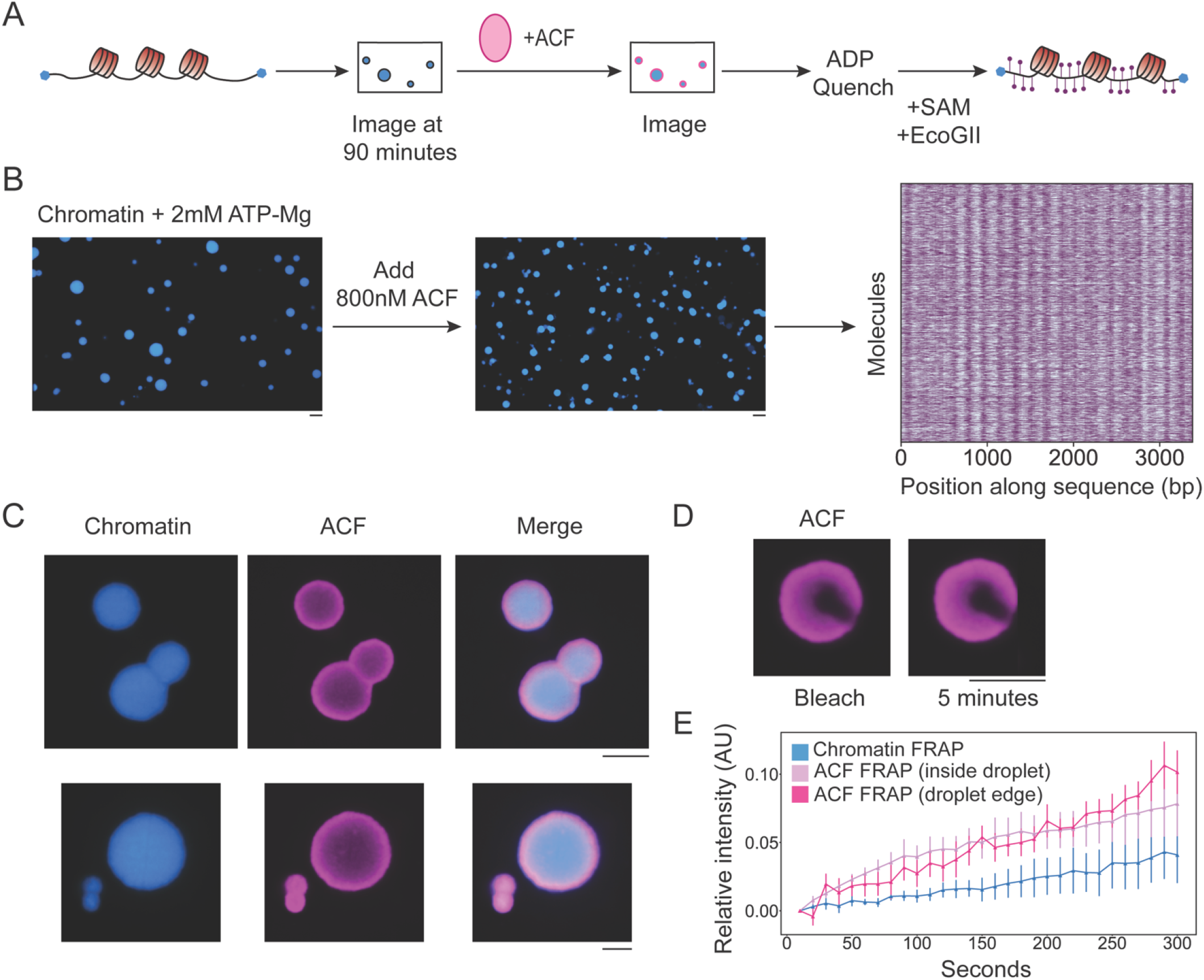
ACF remodels chromatin inside of condensates and is concentrated in condensates. (**A**) Scheme of ACF add-in experimental workflow. (**B**) Alexa647 fluorescence before and after adding in ACF, along with heatmap of accessible and inaccessible bases for 3000 molecules from the add-in reaction. (**C**) Example condensates after adding ACF using AF488-labeled ACF. Chromatin is labeled with AF647. (**D**) Example droplet before and after photobleaching, AF488 channel. (**E**) Quantification of recovery after photobleaching for chromatin (AF647) and ACF (AF488). Recovery is normalized to pre-bleach droplet intensity. All scale bars are 5 um.

To quantify the amount of the ACF within the condensates, we fluorescently labeled ACF and repeated the add-in reaction using condensates generated from arrays with 300nM nucleosome concentrations and 350nM ACF (Fig S4). We observed ACF throughout the condensates, but it was not evenly distributed (Fig 3C, Fig S4E). ACF was approximately two-fold as concentrated (∼12 uM) on the outer surfaces of the chromatin condensates compared to the interior of the condensates (∼6 uM). Based on the mean nucleosome concentration within these condensates of ∼71 uM, we calculated the ratio of ACF:nucleosome inside the condensates as maximally 1:6, and minimally 1:12. This clarified why ACF did not solvate the preformed condensates, because in our previous experiments, ACF detectably inhibited chromatin condensation only when it was near stoichiometric relative to nucleosomes. The decreasing concentration gradient of ACF towards the center of the condensates suggested that ACF was diffusing slowly throughout the condensates. To assay ACF dynamics within the condensates, we FRAPed the labelled ACF. We observed no detectable FRAP at the edge or interior of the condensates over the course of five minutes (Fig 3D-E). It is important to note that our photobleached spots, which are approximately 1μm in diameter, contain tens of thousands of nucleosomes. Thus, the slow recovery does not necessarily imply that ACF is indefinitely stuck on nucleosomes, but rather that it diffuses slower than the order of several minutes across μm-scale distances and through this nucleosome environment. Interestingly, the arrays in these reactions were completely remodeled by ACF at two hours (Fig 3B, S3), indicating that over time, even substochiometric ACF is able to access and remodel all the nucleosomes (Fig S3).

The slow diffusion of ACF through the condensate was not ATP-dependent, as even in the presence of ADP, we observed enrichment of ACF at the periphery of the condensates (Fig S4E). Therefore, we hypothesized that the nM-order K_d_ of ACF for nucleosomes contributed to its slow diffusion through chromatin(*30, 31*). Consistent with this possibility, single molecule FRET data has shown that ACF processively remodels nucleosomes, with ATP-dependent residence times of five minutes or longer on individual mono-nucleosomes(*32*). If high affinity and processivity were responsible for the slow diffusion of ACF through chromatin, then a less processive remodeler with a lower affinity for nucleosomes may diffuse more quickly and uniformly. We therefore repeated the experiment with Snf2h, the catalytic subunit of ACF, which is less processive and has a ∼50-fold weaker affinity for nucleosomes than ACF(*30, 31, 33*). Consistent with our hypothesis, labelled Snf2h recovered from photobleaching much faster, within a minute, and uniformly distributed throughout chromatin condensates in the presence of ATP or ADP (Fig S4 C-D). In congruence with this result, the catalytic subunit of *Drosophila* ISWI complexes, which has a lower affinity for nucleosomes than ACF, diffuses rapidly and uniformly through chromatin condensates within minutes, though only in the presence of ATP (*34*).

Because ACF generates chromatin fibers with regularly spaced nucleosomes, we hypothesized that ACF remodeled products would display more rapid dynamics within the condensates similarly to the evenly spaced 12×601 arrays. However, we observed no FRAP recovery for chromatin remodeled by ACF over 5 minutes (Fig 3E, Fig S4CD). One possible explanation for this result is that even if all nucleosomes are evenly spaced by ACF, the nucleosome spacing varies from molecule to molecule, since some chromatin molecules have more nucleosomes than others. The spacing distributions are best illustrated in the single turnover ACF samples (Fig S3A, 800nM ACF). In contrast, essentially all of the regularly spaced 12×601 array molecules have the same nucleosome spacing due to the strength of this positioning sequence(*28*). It is therefore possible that uniform spacing across all chromatin molecules within a condensate increases dynamics. Another key difference is the presence of ACF, which may make additional interactions that stabilize inter-fiber interactions and slow chromatin dynamics.

In contrast to ACF, RSC generates a wider variety of nucleosomal products so we next investigated whether the different outcomes on nucleosomes result in different outcomes within condensates.

### RSC uses ATP to de-condense chromatin and mobilize whole condensates

We first analyzed how RSC remodeling affected the formation of chromatin condensates by pre-mixing RSC with chromatin containing a median of 14 nucleosomes per template (scheme shown in Fig 4A). In the presence of 100 uM ATP-Mg, we observed that RSC addition resulted in condensates that were smaller and less intense than in the control (Fig 4B, Fig S6). However, RSC inhibited chromatin condensation substantially more in the presence of ADP or AMPPNP (Fig 4B). These results suggested that analogous to ACF, binding by RSC occludes the nucleosomal surfaces that participate in inter-nucleosomal interactions when bound to ADP or AMPNP. However, the differential effect with ATP was not consistent with a simple binding-based model.

**Figure 4.**
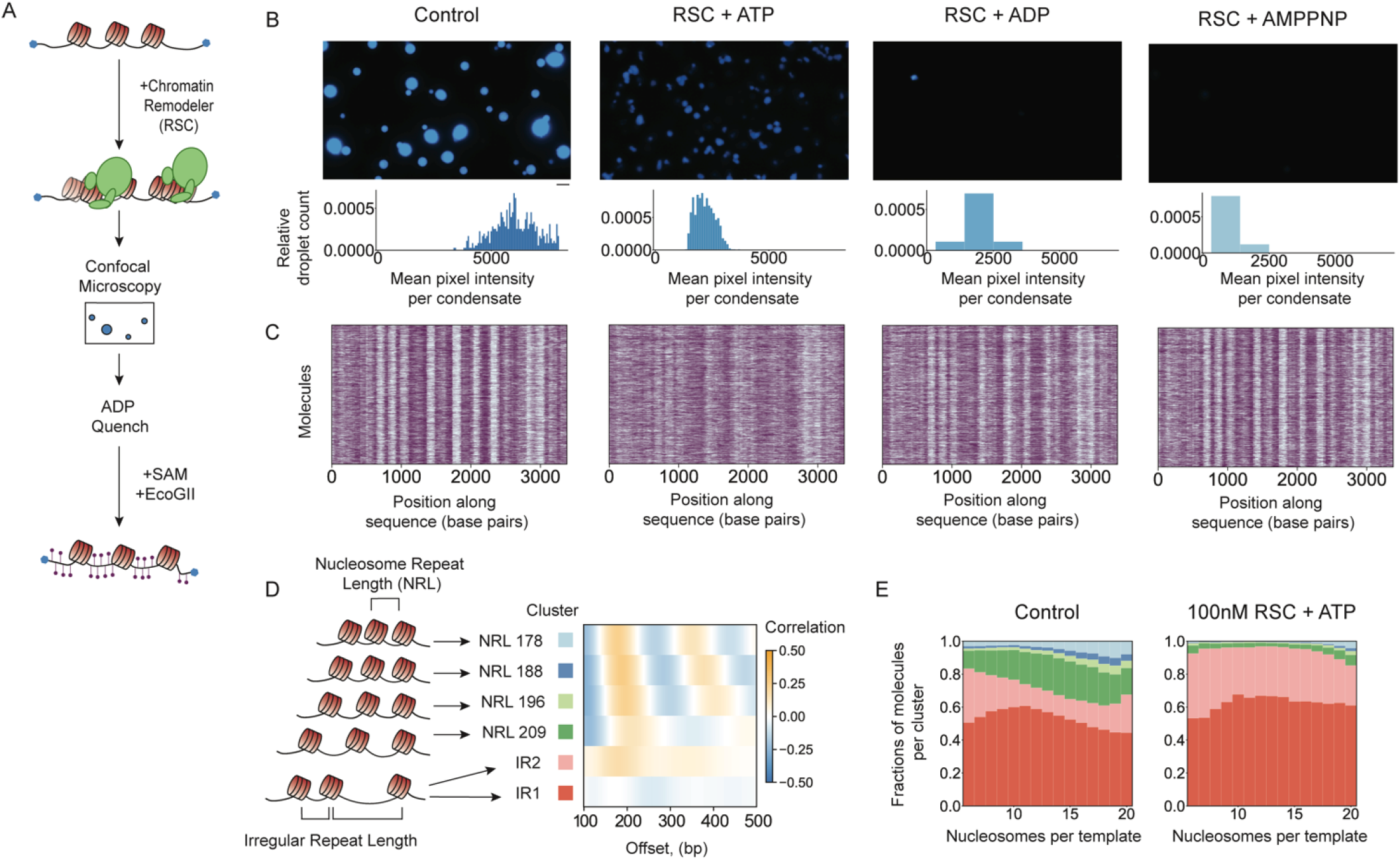
RSC inhibits chromatin condensation and generates diverse nucleosome products that are not evenly spaced. (**A**) Scheme of premix experimental workflow. (**B**) RSC reactions have 100nM RSC, 100uM nucleotide-Mg^2+^, and 150nM nucleosome, with median 15 nucleosomes per molecule. Confocal images show Alexa647 fluorescence and histograms of mean pixel intensity per condensate, with bin size equal to 0.1*number of condensates per condition. Scale bars are 5 um.(C)Heatmaps of accessible and inaccessible bases for 2000 molecules per condition.(D)Average single-molecule autocorrelograms resulting from Leiden clustering of individual molecules. We observe six different array types, four with regular spacing (NRL) and two with irregular spacing (IR). (**E**) Stacked bar chart representation of cluster representation for molecules from two different conditions, plotted as a function of nucleosome density. In the presence of RSC and ATP, the proportion of IR fibers (red and pink colors) increases.

Unlike ACF, the RSC complex slides, disassembles and deforms nucleosomes without evenly spacing them(*5, 12, 35*). These activities are ATP-dependent, but RSC can also bind and distort nucleosomes without ATP(*12, 36*). We observed that RSC substantially displaced nucleosome footprints from their original positions in the presence of ATP (Fig 4C). We also observed minor footprint displacement in the context of ADP (Fig 4C, S5A). In contrast, almost no displacement was seen with AMPPNP (Fig S5A). Interestingly, not all nucleosomes were remodeled to the same extent with ATP, suggesting that the outcome of remodeling depends on DNA sequence (Fig 4C)(*35, 37*). Furthermore, the remodeled nucleosomes were not evenly spaced, as confirmed by autocorrelation analysis (Fig 4D-E). Overall, our analysis indicated that RSC catalyzes outcomes within chromatin condensates are consistent with its previously characterized activities on nucleosomes.

The decreased intensity of condensates in the presence of RSC and ATP suggested that RSC was de-condensing the chromatin. Unexpectedly, we also observed that these condensates moved much more than control droplets (Movie S1-S2). We wondered if preformed chromatin condensates would be similarly de-condensed by RSC. Adding RSC to preformed chromatin condensates made them larger, less intense, and more mobile (Scheme shown in Fig 5A, Fig 5B-E, Movie S3-S6, Fig S6). The volume increase calculated for the condensates accounted for the lower nucleosome concentration (see Methods) consistent with de-condensation rather than loss of chromatin from the droplets. Further, to control for effects of droplet size on droplet motion we size-matched condensates from all four conditions (before RSC and after RSC) and measured the displacement of these condensates over 20 seconds, selecting for condensates that moved > 1.5 um (Fig 5E).

**Figure 5.**
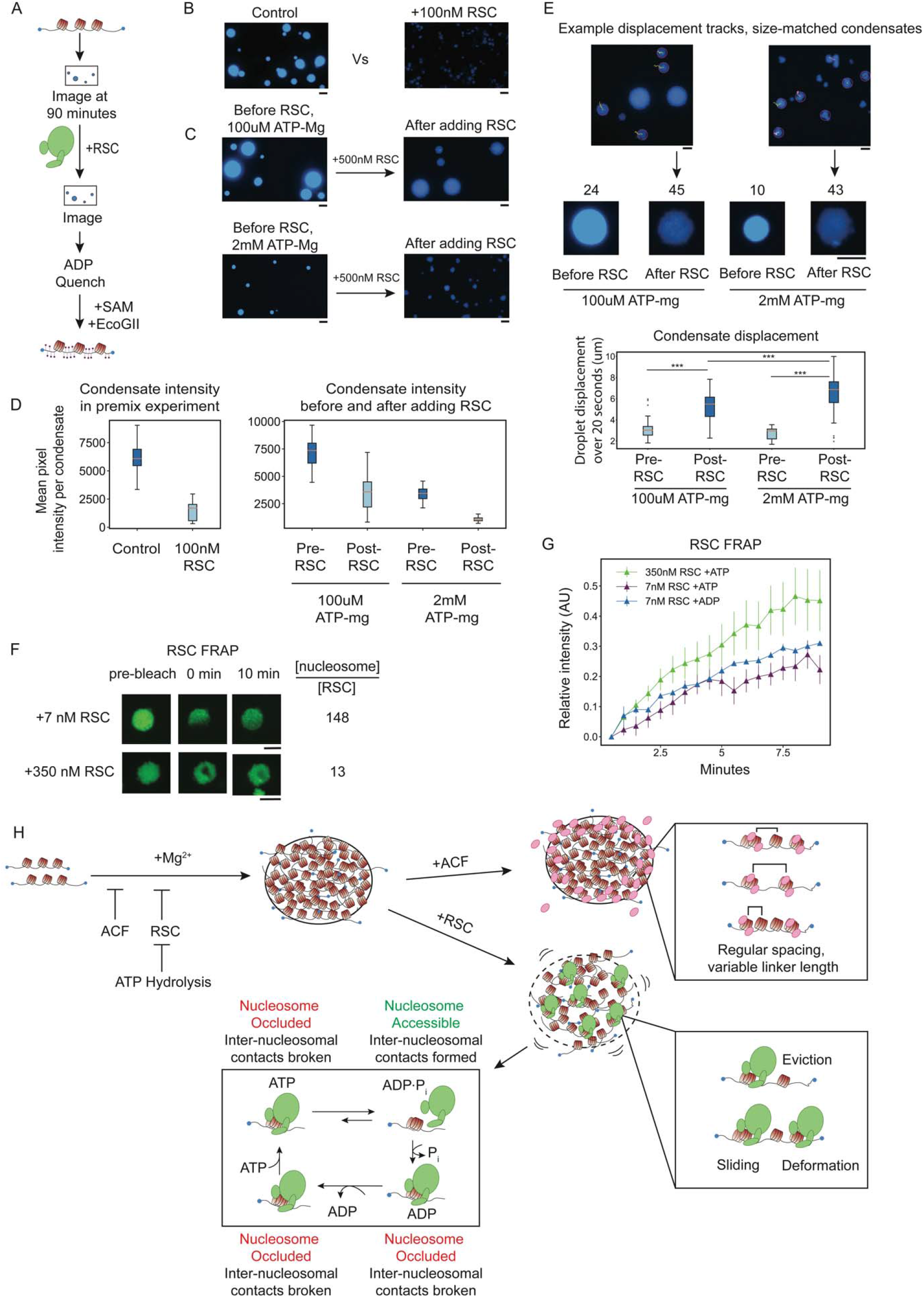
RSC decondenses chromatin condensates and increases condensate motion in an ATP-dependent manner. (**A**) Scheme of RSC add-in experimental workflow. (**B**) Confocal images of chromatin condensates for the RSC premix experiment (scheme shown in Fig 4A) in conditions with 150nM nucleosome, 100uM ATP-Mg. Estimated mean nucleosome concentration inside condensates is 38uM (control reaction) and 9uM (+100nM RSC reaction). (**C**) RSC add-in experiment as shown in A, done in conditions with either 100uM ATP-Mg or 2mM ATP-Mg and 150nM nucleosome. Estimated mean nucleosome concentration inside condensates is 43uM (before RSC, 100uM ATP-mg), 21uM (after RSC, 100uM ATP-mg), 21uM (before RSC, 2mM ATP-mg), 7uM (after RSC, 2mM ATP-mg). (**D**) Boxplots of mean condensate intensity for premix (left) and add in experiments in A and B (right). (**E**) Example traces of condensate displacement over 20 seconds. Displacement analysis was limited to condensates with diameter of approximately 5 um and the number of 5 um diameter condensates tracked in each condition is listed above the example droplets. (**E**) Boxplot of size-matched condensate displacement before and after RSC add-in. (**F**) On left, confocal images of RSC add-in using AF488 labeled RSC, 488 channel. Top row images are from 7nM AF488-RSC add-in, bottom row images from 350nM AF488-RSC add-in. Both were done in conditions with 100uM ATP-Mg. The molar ratio of nucleosome to RSC inside condensates is shown. (**G**) AF488 fluorescence recovery after photobleaching for 7nM and 350nM RSC. Recovery is normalized to pre-bleach droplet intensity and plotted an average of 6 droplets for 350nM RSC and 3 droplets for 7nM RSC. (**H**) Model for chromatin condensation and remodeling within the condensate by ACF or RSC. All scale bars are 5 um.

The increased droplet motion and chromatin de-condensation seen with RSC could be a consequence of the remodeled chromatin products or could require active ATP hydrolysis. To uncouple the effects of remodeling outcome from active ATP hydrolysis, we compared RSC action at 100 uM and 2 mM ATP. At both of these ATP concentrations, the chromatin is completely remodeled at the time of imaging. Yet, there was more condensate motion at the higher ATP concentration (Movie S3-6). These results indicate that the condensate motion observed after adding RSC and ATP depends on active ATP hydrolysis.

These results uncover a major difference between how ACF and RSC affect compacted chromatin. Unlike with ACF, RSC has two ATP-hydrolysis dependent effects: (1) decompaction of chromatin condensates without dissolution and (2) increase of condensate motion. To better understand how RSC may drive condensate motion we investigated how RSC was distributed within the condensates (Fig. S5). Unlike ACF, we found that RSC is evenly distributed throughout condensates with ATP or ADP and recovers from photobleaching within 10 minutes (Fig 5F-G). However similar to ACF, we observed sub-stoichiometric RSC relative to nucleosomes (1 RSC:13 nucleosomes) inside the condensates even when stoichiometric concentrations of RSC were added into the condensates (Fig 5F). As elaborated in the discussion, we propose that the larger size of RSC and its lower processivity compared to ACF may explain its unique ATP-dependent effects on condensates.

## Discussion

Nucleosome concentrations are estimated to range from ∼100-500uM in vivo (*22*), and it is poorly understood how chromatin remodelers act within such crowded environments. Chromatin condensates allow recreation of comparably crowded conditions *in vitro*. Here, by carrying out mechanistic biochemistry in chromatin condensates, we have uncovered new consequences of ATP-dependent chromatin remodeling by ACF and RSC, two remodelers that respectively enable heterochromatin and euchromatin formation. Below we discuss the biological implications of our findings.

### Nucleosome spacing and density regulate the properties of condensed chromatin

Chromatin condensation depends on inter-nucleosomal interactions. Therefore, nucleosome density is expected to influence the formation and compaction of condensates. Here, we systematically tested this prediction. Keeping DNA concentrations constant, we found that median densities of 4.6 nucleosome/kb yield condensates with nucleosome concentrations comparable to *in vivo*, densities of 2.9 nucleosome/kb yield condensates with a fivefold lower concentration of nucleosomes, and densities of 1 nucleosome/kb do not yield detectable condensates. Furthermore, condensates formed by chromatin assembled on the native S3 sequence show increased viscosity relative to condensates from chromatin with regularly spaced nucleosomes at comparable density. Recent Cryo-ET studies of native mammalian chromatin show a high proportion of irregularly spaced nucleosomes and find that such chromatin is found in short intra-fiber stacks of nucleosomes in cis interspersed with inter-fiber interactions in trans (*38*). In contrast, regularly spaced nucleosome arrays appear to contain a greater proportion of intra-fiber nucleosome stacks (*38-40*). These differences may result in more stable inter-fiber interactions with irregularly spaced nucleosomes, explaining the greater viscosity.

### Remodeler affinity and processivity govern diffusion within condensed chromatin

Although the chromatin condensates are viscous, nucleosomes within the condensates are still accessible to nucleosome remodelers. Both ACF and RSC can diffuse throughout chromatin condensates and remodel the nucleosomes within. Interestingly, diffusion of ACF is much more restricted than that of RSC. We propose that this difference arises from the differences in the kinetics of how ACF and RSC engage nucleosomes. ACF has been shown to move nucleosomes in a bidirectional and highly processive manner, remaining bound to nucleosomes for several minutes during a remodeling cycle(*32*). In contrast, the residence time of RSC on nucleosomes is on the order of seconds(*41*). Therefore, within a condensate, ACF would be trapped on a nucleosome more often, slowing its effective diffusion relative to RSC. Consistent with this hypothesis, we find that the catalytic subunit Snf2h, which has lower nucleosome affinity and lower processivity, diffuses more rapidly throughout the chromatin condensates.

Chromatin condensates also seem to inherently limit remodeler concentrations. Both ACF and RSC, when added to pre-formed condensates in stoichiometric concentrations to nucleosomes, only achieve sub-stoichiometric enrichment within condensates. We speculate that the network of inter-nucleosomal interactions within condensates limits the number of remodeler molecules that can be accommodated. Thus, diffusion of a remodeler through a chromatin condensate would be regulated by both the off rate of the remodeler and the kinetics of breaking the local network of inter-nucleosomal interactions. Interestingly, single particle tracking studies in yeast show much faster diffusion of remodelers through the yeast nucleus than we observe in our reconstituted condensates (*42*). It is possible that in cells, additional factors compete with remodelers to drive facilitated dissociation or help disrupt inter-nucleosomal interactions (*43*).

ACF has been shown to play a critical role in maintaining heterochromatin domains through its nucleosome spacing activity. Our findings suggest that ACF’s high affinity and processivity may allow for its sustained localization within condensed heterochromatin. By comparison, the faster diffusion by RSC may enable faster decondensation of a chromatin domain during the formation of euchromatin and increase the local fluidity of chromatin as discussed below.

### Nucleosome scale remodeling can explain meso-scale consequences

We find that ACF and RSC have different effects within condensed chromatin. We propose that these differences arise from two features (1) how ACF and RSC bind and remodel nucleosomes (Fig 5H) and (2) their residency times on nucleosomes during ATP-hydrolysis. Compared to ACF, RSC, which is bigger, is expected to occlude a larger region of the nucleosome and therefore have a larger effect on disrupting internucleosomal interactions (*44-46*). However, during ATP hydrolysis, the residency time of RSC on nucleosomes has been shown to be shorter than for ACF (*47*). We propose that ATP hydrolysis switches RSC between nucleosome-bound states that inhibit inter-nucleosomal interactions and nucleosome free states that allow inter-nucleosomal interactions (Fig 5H). We speculate that such ATP-driven cycling between states transiently and locally disrupts inter-nucleosomal interactions without globally dissolving the condensate. Such a cycle may increase the local dynamics within condensed chromatin. Interestingly, RSC catalyzes whole condensate motion despite being substantially sub-stoichiometric relative to nucleosomes (Fig 5H). This result suggests that disrupting some inter-nucleosomal interactions can have cooperative effects.

*In vivo*, remodelers likely need to engage nucleosomes in at least two different contexts: in open chromatin states prior to condensation, and in states where the chromatin is condensed. ACF has been implicated in regulating heterochromatin(*8, 48*). We speculate that newly replicating heterochromatin would be prevented from prematurely condensing by stoichiometric ACF binding and remodeling. In contrast, in the context of an existing heterochromatin domain, our findings imply that ACF can maintain the nucleosome spacing within heterochromatin without decondensing chromatin. Our studies also clarify that ACF does not require other nuclear proteins in order to penetrate, remodel, and remain localized at condensed chromatin domains. In contrast to ACF, we find that RSC decondenses pre-formed chromatin condensates in an ATP-hydrolysis dependent manner. It is well established that chromatin de-condensation correlates with transcription, but genomic studies cannot evaluate the effects of RSC on chromatin condensation in a transcription-independent manner, because RSC mutations affect transcription(*16, 49*). Our approach reveals that RSC remodeling does induce transcription-independent chromatin de-condensation, which likely contributes to de-condensation of active gene promoters *in vivo*. Additionally, we speculate that the increased condensate motion caused by RSC may enable faster local diffusion of the transcription machinery and other proteins through the viscous environment of condensed chromatin.

We demonstrate that chromatin remodelers impact chromatin beyond the scale of individual nucleosomes, influencing the accessibility and dynamics of kilobase-size chromatin domains. Our findings further demonstrate that ATP-dependent remodelers do not generically act as molecular stir bars; rather, their meso-scale effects on chromatin derive from their specific, nucleosome-scale interactions and activities. Future work is needed to clarify whether other nucleosome remodelers that catalyze distinct transformations of nucleosomes also have unique effects on meso-scale chromatin dynamics.

## Acknowledgements

We thank J. Tretyakova for histone purification, L. Hsieh for RSC construct and invaluable advice, K. Harrington and S.Y. Kim for their expansive knowledge of the confocal microscope, M. Rosen (UT Southwestern) for the 12×601 plasmid, S. Nanda for mentorship, and all members of the Narlikar and Ramani labs for advice and discussion. We also thank Bo Huang for noticing the movement of droplets with RSC and encouraging us to pursue it.

## Author Contributions

C.M. and U.K. purified RSC; C.M. and Z.Z. purified ACF; U.K. and U.S.C. assembled and purified mono-nucleosomes; E.W. purified 12×Widom601 DNA; M.O., K.W., S.W., I.I., and C.M. prepared samples for PacBio sequencing; C.M. performed and quantified all biochemical experiments; V.R. and G.J.N. conceived and oversaw the project; C.M., V.R., and G.J.N. participated in interpretation and discussion of the results and writing of the manuscript.

## Data and materials availability

Raw and processed data will be made available at GEO. All scripts and notebooks used for data analysis will be available on Github.

## Supplementary Materials

Materials and Methods

Figs. S1 to S7

References(50-55)

Movies S1 to S6

## References

1. E. Lieberman-Aiden et al., Comprehensive Mapping of Long-Range Interactions Reveals Folding Principles of the Human Genome. Science 326, 289–293 (2009).

2. J. Dekker, Mapping in vivo chromatin interactions in yeast suggests an extended chromatin fiber with regional variation in compaction. J Biol Chem 283, 34532–34540 (2008).

3. C. R. Clapier, J. Iwasa, B. R. Cairns, C. L. Peterson, Mechanisms of action and regulation of ATP-dependent chromatin-remodelling complexes. Nature Reviews Molecular Cell Biology 18, 407–422 (2017).

4. C. Y. Zhou, S. L. Johnson, N. I. Gamarra, G. J. Narlikar, Mechanisms of ATP-Dependent Chromatin Remodeling Motors. Annu Rev Biophys 45, 153–181 (2016).

5. N. Krietenstein et al., Genomic Nucleosome Organization Reconstituted with Pure Proteins. Cell 167, 709-721.e712 (2016).

6. K. Luger, A. W. Mäder, R. K. Richmond, D. F. Sargent, T. J. Richmond, Crystal structure of the nucleosome core particle at 2.8 Å resolution. Nature 389, 251–260 (1997).

7. T. Ito, M. Bulger, M. J. Pazin, R. Kobayashi, J. T. Kadonaga, ACF, an ISWI-Containing and ATP-Utilizing Chromatin Assembly and Remodeling Factor. Cell 90, 145–155 (1997).

8. D. V. Fyodorov, M. D. Blower, G. H. Karpen, J. T. Kadonaga, Acf1 confers unique activities to ACF/CHRAC and promotes the formation rather than disruption of chromatin in vivo. Genes & development 18, 170–183 (2004).

9. T. Tsukiyama, J. Palmer, C. C. Landel, J. Shiloach, C. Wu, Characterization of the imitation switch subfamily of ATP-dependent chromatin-remodeling factors in Saccharomyces cerevisiae. Genes Dev 13, 686–697 (1999).

10. P. D. Varga-Weisz et al., Chromatin-remodelling factor CHRAC contains the ATPases ISWI and topoisomerase II. Nature 388, 598–602 (1997).

11. G. Längst, E. J. Bonte, D. F. V. Corona, P. B. Becker, Nucleosome Movement by CHRAC and ISWI without Disruption or trans-Displacement of the Histone Octamer. Cell 97, 843–852 (1999).

12. Y. Lorch, M. Zhang, R. D. Kornberg, RSC Unravels the Nucleosome. Molecular Cell 7, 89–95 (2001).

13. S. Ramachandran, G. E. Zentner, S. Henikoff, Asymmetric nucleosomes flank promoters in the budding yeast genome. Genome research 25, 381–390 (2015).

14. B. R. Cairns et al., RSC, an essential, abundant chromatin-remodeling complex. Cell 87, 1249–1260 (1996).

15. M. Floer et al., A RSC/Nucleosome Complex Determines Chromatin Architecture and Facilitates Activator Binding. Cell 141, 407–418 (2010).

16. T. H. Hsieh et al., Mapping Nucleosome Resolution Chromosome Folding in Yeast by Micro-C. Cell 162, 108–119 (2015).

17. K. Yen, V. Vinayachandran, K. Batta, R. T. Koerber, B. F. Pugh, Genome-wide nucleosome specificity and directionality of chromatin remodelers. Cell 149, 1461–1473 (2012).

18. A. Klein-Brill, D. Joseph-Strauss, A. Appleboim, N. Friedman, Dynamics of Chromatin and Transcription during Transient Depletion of the RSC Chromatin Remodeling Complex. Cell Rep 26, 279-292.e275 (2019).

19. S. Kubik et al., Sequence-Directed Action of RSC Remodeler and General Regulatory Factors Modulates +1 Nucleosome Position to Facilitate Transcription. Mol Cell 71, 89-102.e105 (2018).

20. E. Oberbeckmann, K. Quililan, P. Cramer, A. M. Oudelaar, In vitro reconstitution of chromatin domains shows a role for nucleosome positioning in 3D genome organization. Nat Genet 56, 483–492 (2024).

21. A. G. Larson, G. J. Narlikar, The Role of Phase Separation in Heterochromatin Formation, Function, and Regulation. Biochemistry 57, 2540–2548 (2018).

22. S. Hihara et al., Local Nucleosome Dynamics Facilitate Chromatin Accessibility in Living Mammalian Cells. Cell Reports 2, 1645–1656 (2012).

23. B. A. Gibson et al., Organization of chromatin by intrinsic and regulated phase separation. Cell 179, 470-484. e421 (2019).

24. J. C. Hansen, K. Maeshima, M. J. Hendzel, The solid and liquid states of chromatin. Epigenetics & Chromatin 14, 50 (2021).

25. N. J. Abdulhay et al., Nucleosome density shapes kilobase-scale regulation by a mammalian chromatin remodeler. Nature Structural & Molecular Biology 30, 1571–1581 (2023).

26. B. A. Gibson et al., In diverse conditions, intrinsic chromatin condensates have liquid-like material properties. Proceedings of the National Academy of Sciences 120, e2218085120 (2023).

27. K. Maeshima et al., A Transient Rise in Free Mg2+ Ions Released from ATP-Mg Hydrolysis Contributes to Mitotic Chromosome Condensation. Current Biology 28, 444-451.e446 (2018).

28. N. J. Abdulhay et al., Massively multiplex single-molecule oligonucleosome footprinting. eLife 9, e59404 (2020).

29. A. S. Nanda et al., Direct transposition of native DNA for sensitive multimodal single-molecule sequencing. Nature Genetics, (2024).

30. J. G. Yang, T. S. Madrid, E. Sevastopoulos, G. J. Narlikar, The chromatin-remodeling enzyme ACF is an ATP-dependent DNA length sensor that regulates nucleosome spacing. Nature Structural & Molecular Biology 13, 1078–1083 (2006).

31. X. He, H. Y. Fan, G. J. Narlikar, R. E. Kingston, Human ACF1 alters the remodeling strategy of SNF2h. J Biol Chem 281, 28636–28647 (2006).

32. T. R. Blosser, J. G. Yang, M. D. Stone, G. J. Narlikar, X. Zhuang, Dynamics of nucleosome remodelling by individual ACF complexes. Nature 462, 1022–1027 (2009).

33. S. L. Johnson, G. J. Narlikar, ATP Hydrolysis Coordinates the Activities of Two Motors in a Dimeric Chromatin Remodeling Enzyme. J Mol Biol 434, 167653 (2022).

34. P. Vizjak et al., ISWI catalyzes nucleosome sliding in condensed nucleosome arrays. Nature Structural & Molecular Biology, (2024).

35. C. E. Rowe, G. J. Narlikar, The ATP-dependent remodeler RSC transfers histone dimers and octamers through the rapid formation of an unstable encounter intermediate. Biochemistry 49, 9882–9890 (2010).

36. B. T. Harada et al., Stepwise nucleosome translocation by RSC remodeling complexes. eLife 5, e10051 (2016).

37. P. D. Partensky, G. J. Narlikar, Chromatin remodelers act globally, sequence positions nucleosomes locally. J Mol Biol 391, 12–25 (2009).

38. N. Jentink, C. Purnell, B. Kable, M. T. Swulius, S. A. Grigoryev, Cryoelectron tomography reveals the multiplex anatomy of condensed native chromatin and its unfolding by histone citrullination. Molecular Cell 83, 3236-3252.e3237 (2023).

39. B. Dorigo et al., Nucleosome arrays reveal the two-start organization of the chromatin fiber. Science 306, 1571–1573 (2004).

40. A. Routh, S. Sandin, D. Rhodes, Nucleosome repeat length and linker histone stoichiometry determine chromatin fiber structure. Proc Natl Acad Sci U S A 105, 8872–8877 (2008).

41. J. M. Kim et al. (Cold Spring Harbor Laboratory, 2023).

42. J. M. Kim et al., Single-molecule imaging of chromatin remodelers reveals role of ATPase in promoting fast kinetics of target search and dissociation from chromatin. eLife 10, e69387 (2021).

43. R. I. Kamar et al., Facilitated dissociation of transcription factors from single DNA binding sites. Proc Natl Acad Sci U S A 114, E3251–e3257 (2017).

44. A. B. Patel et al., Architecture of the chromatin remodeler RSC and insights into its nucleosome engagement. eLife 8, e54449 (2019).

45. F. R. Wagner et al., Structure of SWI/SNF chromatin remodeller RSC bound to a nucleosome. Nature 579, 448–451 (2020).

46. L. Li et al., Structure of the ISW1a complex bound to the dinucleosome. Nat Struct Mol Biol 31, 266–274 (2024).

47. J. M. Kim et al., Dynamic 1D search and processive nucleosome translocations by RSC and ISW2 chromatin remodelers. eLife 12, RP91433 (2024).

48. N. Collins et al., An ACF1–ISWI chromatin-remodeling complex is required for DNA replication through heterochromatin. Nature Genetics 32, 627–632 (2002).

49. L. Lopez-Serra, G. Kelly, H. Patel, A. Stewart, F. Uhlmann, The Scc2-Scc4 complex acts in sister chromatid cohesion and transcriptional regulation by maintaining nucleosome-free regions. Nat Genet 46, 1147–1151 (2014).

50. K. Luger, T. J. Rechsteiner, T. J. Richmond, Expression and purification of recombinant histones and nucleosome reconstitution. Methods Mol Biol 119, 1–16 (1999).

51. N. Gamarra, S. L. Johnson, M. J. Trnka, A. L. Burlingame, G. J. Narlikar, The nucleosomal acidic patch relieves auto-inhibition by the ISWI remodeler SNF2h. eLife 7, e35322 (2018).

52. C. Y. Zhou et al., The Yeast INO80 Complex Operates as a Tunable DNA Length-Sensitive Switch to Regulate Nucleosome Sliding. Mol Cell 69, 677-688.e679 (2018).

53. C. A. Schneider, W. S. Rasband, K. W. Eliceiri, NIH Image to ImageJ: 25 years of image analysis. Nature Methods 9, 671–675 (2012).

54. D. Ershov et al., TrackMate 7: integrating state-of-the-art segmentation algorithms into tracking pipelines. Nature Methods 19, 829–832 (2022).

55. H. Li, Minimap2: pairwise alignment for nucleotide sequences. Bioinformatics 34, 3094–3100 (2018).

