## Supplemental Files for "ATP-dependent remodeling of chromatin condensates uncovers distinct mesoscale effects of two remodelers"

**The PDF file includes:**

Materials and Methods

Figs. S1 to S7

References (50-55)

**Other Supplementary Materials for this manuscript include the following:**

Movies S1 to S12

Materials and Methods

**Construction of *Cyp3a11* dsDNA-producing Bacterial Vector**

A 3.5 kilobase stretch of Cyp3a11 was PCR amplified from mouse genomic DNA. This fragment was cloned into the puc18 plasmid using Gibson Cloning (NEBuilder HiFi DNA Assembly Master Mix, NEB) and transformed into DCM-/DAM- competent cells. Colonies were isolated and miniprepped (Qiagen) and plasmid sequence was validated via Primordium Sequencing.

**Expression and Purification of Recombinant Proteins**

Recombinant histones from X. laevis were expressed and purified in E. coli as previously described(*50*). The Snf2h ATPase was purified from *E. coli* (*51*) and the human ACF complex was purified from Sf9 insect cells as previously described(*51*) with a minor modification. ACF1-FLAG and SNF2H were expressed on the same plasmid via infection with baculovirus.

The RSC2 C-terminally 3xFLAG-tagged yeast strain was generated by Elise Munoz and Laura Hsieh. The FLAG-tagged RSC complex was purified from yeast as previously described(*52*).

Protein concentrations were determined via SYPRO red (Thermo Fisher) staining of a sodium dodecyl sulfate (SDS)–polyacrylamide gel electrophoresis gel with bovine serum albumin standards.

**Array DNA Purification**

12x601+47 plasmid was a generous gift from the Rosen lab. The 12x601 plasmid was transformed into Stable 3 cells and purified via Giga Prep (Qiagen). The 12x601 insert was isolated from the plasmid backbone via restriction digest (EcoRV-HF) and size exclusion chromatography. After size exclusion, purified 12x601 insert was precipitated in ethanol and resuspended in 1X TE.

The *Cyp3a11* sequence was digested from the puc18 backbone via PciI and BamHI and similarly purified.

**DNA Labeling**

Cypa11 array DNA was end-labeled with Alexa Fluor-647-aha-dCTP or Alexa Fluor-555-aha-dCTP (Thermo) using the Klenow fragment.

12x601 array DNA was first digested with XhoI to generate a 5’ overhang, then similarly labeled with Alexa Fluor-647-aha-dCTP using the Klenow fragment. Labeling efficiency was quantified via NanoDrop.

**Histone Octamer Purification**

Histones were refolded in high salt buffer mixed to form octamer. Octamer was purified by size-exclusion chromatography as previously described(*50*).

**Chromatin Assembly**

Chromatin was assembled using salt gradient dialysis with varying ratios of histone octamer:DNA(*50*). DNA concentration was determined after assembly via Nanodrop.

**Sucrose Gradient**

After salt gradient dialysis, 12x601 chromatin arrays were added to a 10-30% sucrose gradient (10mM Tris pH 7.5, 1mM EDTA, 1mM DTT, sucrose) and spun for 16 hours at 22,900 RPM. Fractions containing assembled nucleosomes were concentrated using 10,000 MWCO centrifugal concentrators (Amicon).

**Protein Labeling**

Remodelers were labeled with Alexa Fluor-488 C_5_ Maleimide (Thermo). Purified remodeler was dialyzed overnight to remove DTT, then reduced via 10-fold molar excess of TCEP relative to the protein. Dye was resuspended in DMSO. Labeling was done with 20-fold molar excess dye relative to the protein for 40 minutes at room temperature, then quenched with excess DTT. Free dye was removed from labeled protein via overnight dialysis in 500mL buffer using a pump to continuously flow buffer (1L total buffer was flowed through). Buffer compositions were identical to original buffer compositions before labeling. Protein concentration was calculated via Nanodrop and SYPRO red (Thermo Fisher) staining of SDS–polyacrylamide gel electrophoresis gel with bovine serum albumin standards. Labeling efficiency was calculated via Nanodrop.

**Mono-nucleosome Remodeling Assay**

Mono-nucleosomes were prepared via salt gradient dialysis from purified X. laevis histone octamers and PCR-purified Widom601 sequence + 80 bp or Cy5-40bp-Widom601 sequence-40 bp as previously described**.** Mono-nucleosomes were remodeled via ACF, Snf2h, and RSC under single turnover saturating enzyme and saturating ATP conditions. Reactions contained 30mM HEPES pH 7.5, 50mM KCl, 1mM MgCl_2, ._0.02% NP-40, 5mM ATP-MgCl_2_. Time points taken from the reaction were quenched with 0.4mg/ml puc19, 20mM ADP, 8% glycerol. Nucleosomes were loaded on a 6% polyacrylamide gel in 0.5x TBE and run at 150V for 2 hours. DNA bands were imaged using the Cy5 channel or SybrSafe stain on a Typhoon imager (GE Life Sciences).

**Preparation of Microscopy Plates**

Corning 384-well microscopy plates were mPEGylated and passivated with bovine serum albumin (BSA) as previously described(*23*).

**Phase Separation Remodeling Reactions**

Unless otherwise stated, phase separation reactions contained 25nM chromatin array, 100mM KCl, 7% glycerol, 25mM HEPES pH 7.5, 5mM DTT, 2mM free Mg2+, 0.5mM EDTA. ATP-Mg concentration was either 100uM or 2mM.

Phase separation reactions in Figure 1 contained 75mM KCl, 2.5% glycerol, 20mM HEPES pH 7.5, 2mM ATP, 4 mM Mg2+, 0.5mM EDTA.

Phase separation reaction conditions for the 12x601 array were 150mM NaCl, 5% glycerol, 25mM Tris-Cl pH 7.5, 1mM MgCl_2_, 5mM DTT.

Reactions were mixed and added to a PEGylated and BSA-passivated microscopy plate. Reactions were sealed with PCR foil to prevent evaporation. After 1 hour, foil was removed and reactions were imaged using a spinning disk confocal microscope. For add-in reactions, protein was added after initial imaging and gently pipetted several times to mix. 30 minutes after mixing, add-in reactions were imaged again and FRAPped.

After imaging, excess ADP (34mM final) was added to reactions and mixed with a pipette several times to quench ATP-dependent remodeling.

**Microscopy**

Data for this study were acquired at the Center for Advanced Light Microscopy at UCSF. Confocal microscopy images were acquired using a Nikon Ti Eclipse microscope base equipped with either a Yokogawa CSU-22 spinning disk confocal unit or CREST X-Light V2 L-FOV Spinning Disk confocal unit, 100 X 1.40 NA oil objective, and an Andor Zyla 4.2 camera. Fluorescence Recovery After Photobleaching (FRAP) was done with a 473 nm laser (Vortran) and Rapp UGA-40 photobleaching system. Widefield microscopy images in Figure 2D were acquired using a Nikon Ti Eclipse microscope base equipped with a 20 x 0.75 NA objective and a Nikon DS-Qi2 camera.

**Image Intensity and Displacement Quantification**

Image analysis was done with ImageJ (Version 2.14)(*53*). Unless otherwise described, image brightness and contrast were normalized for a given panel of images and images were collected under identical microscopy settings. Mean pixel intensities per chromatin condensate were calculated in ImageJ from .tif files. Condensates were picked using the Otsu algorithm and auto thresholding using minimum cutoffs of 0.2 circularity and 0.5 um^2^. Condensates on image edges were excluded from analysis. Displacement tracking analysis of chromatin droplets was done with the Trackmate plugin in ImageJ(*54*). In Figure 5, minimum droplet displacement was set to 1.5 um over 20 seconds.

**Correlating chromatin condensate area change with condensate intensity change after adding RSC**

The fold change in condensate area was computed by dividing the mean condensate area after adding RSC by the mean condensate area before adding RSC (area distributions shown in Fig S6A). For the low ATP conditions (100uM ATP-mg), the fold change in area was 44.64/24.08=1.85. For the high ATP condition (2mM ATP-mg), the fold change in area was 8.35/3.67 = 2.28.

Fold change in volume = (fold change in area)^3/2^

Fold change in volume after adding RSC in conditions with 100uM ATP-mg: 2.5

Fold change in volume after adding RSC in conditions with 2mM ATP-mg: 3.4

This ratio should be proportional to the decrease in mean pixel intensity per condensate after adding RSC if chromatin is maintained within the condensates after adding RSC.

The decrease in pixel intensity per condensate for the AF647 channel was computed by dividing the mean pixel intensity per condensate before adding RSC by the mean pixel intensity per condensate after adding RSC (pixel intensity distributions shown in Fig 5D).

Fold change in pixel intensity per condensate after adding RSC in in conditions with 100uM ATP-mg: 2.08

Fold change in pixel intensity per condensate after adding RSC in in conditions with 2mM ATP-mg: 3.15

**Calculating nucleosome concentration inside condensates**

Nucleosome concentration was determined using the mean AF647 channel pixel intensity per condensate and a standard curve of free AF647 dye using the same exposure and laser power as samples (Fig S1). The mean pixel intensity per condensate was divided by two because each chromatin molecule has two fluorescent labels (Fig 1A), and this value was divided by the slope determined by the standard curve (1235.5/uM), then multiplied by the median number of nucleosomes per molecule determined for that chromatin assembly.

Condensate nucleosome concentration = (mean pixel intensity per condensate)/2 * (1235.5/uM) * X nucleosomes/molecule

**Calculating remodeler concentration inside condensates**

Remodeler concentrations were determined using the mean AF488 channel pixel intensity per condensate and a standard curve of free AF488 dye using the same exposure and laser power as samples (Fig S1). The mean pixel intensity per condensate was divided by the molar ratio of label/protein determined via Nanodrop, and this value was divided by the slope determined using the AF488 standard curve.

**SAMOSA-ChAAT on chromatin arrays**

SAMOSA-ChAAT was performed on chromatin arrays using the nonspecific adenine methyltransferase EcoGII (NEB, high concentration stock 2.5 x 10^4^ U ml^-1^) as previously described(*25, 28*) with minor modifications. The phase separation reaction volume was removed from the microscopy plate and diluted to 100 ul in 1xCutSmart buffer + 1 mM SAM + 1ul EcoGII. Reactions were mixed and incubated at 37C for 30 minutes. After 30 minutes, 10ul of 10% SDS and 2.5 ul Proteinase K (20 mg/ml) was added to each reaction and mixed with a pipette. Reactions were incubated at 65C for 2 hours or overnight. Methylated DNA was purified from these reactions via 1X SPRI Select Beads.

**Pacbio Library Preparation and Sequencing**

Fluorophores were removed from DNA by restriction digest in 50 ul of 1x CutSmart with 1ul SmaI and 1 ul BsiEI. Restriction digests were incubated for 15 minutes at room temperature, then 15 minutes at 60C. DNA was purified from these reactions via 1X SPRI Select Beads.

Entire remodeling reactions were used as input for PacBio SMRTbell library preparation. SMRTbell preparation of libraries was done using the SMRTbell prep kit 3.0 and included DNA damage repair, end repair, SMRTbell ligation, and exonuclease cleanup according to the manufacturer’s instruction. After exonuclease cleanup and purification via 1x v/v SMRTbell cleanup beads, DNA concentration was measured by Qubit High Sensitivity DNA Assay (1 ul each sample). Data was collected over 30-hour Sequel II movie runs with 2 hours pre-extension time and 2.1 polymerase.

**SMRT Data Processing**

Sequencing reads were processed as homogenous samples as described in (*25*) with slight variations.

**Model Training**

For training neural network, SMM, and SVD models on fully methylated and unmethylated controls, raw subreads were processed identically to homogenous samples(*25*) and models were trained as previously described. The Hidden Markov model was structured similarly to (*25*) but was refactored from pomegranate to use cython and numba.

**Chromatin Sample Processing**

Raw sequencing reads from chromatin samples were processed using software from Pacific Biosciences:

1. Generate circular consensus sequences (CCS)

CCS were generated for each sequencing cell using ccs 6.9.99. The --hifi-kinetics flag was used to generate kinetics information (interpulse duration, or IPD) for each base of each consensus read. Values were stored for each base as 50*(mean logIPD) + 1.

2. Demultiplex consensus reads

Consensus reads were demultiplexed using lima. The flag ‘–same’ was passed as libraries were generated with the same barcode on both ends. This produces a BAM file for the consensus reads of each sample.

3. Align consensus reads to the reference genome

pbmm2, the pacbio wrapper for minimap2(*55*), was run on each CCS BAM file (the output of step 2) to align reads to the reference sequence, producing a BAM file of aligned consensus reads.

**Extracting interpulse duration measurements**

The IPD values were accessed from the aligned, demultiplexed consensus BAM files. Values were transformed so that each value represented the log_10_IPD, in order to match the log_10_IPD values that were used to train the models.

**Processed data analysis**

All processed data analyses and associated scripts are available at GitHub. All analyses were computed using python. Plots were constructed via Matplotlib. Each analysis is briefly described below:

**Defining inaccessible regions and counting nucleosomes**

Inaccessible regions were called from HMM output data identically to (*25*). Briefly, inaccessible regions were defined as continuous stretches with accessibility ≤0.5. Periodic peaks were observed that approximated sizes of regions containing one, two, three, or more nucleosomes. Cutoffs for each size were manually defined using the histogram of inaccessible region lengths (Figure S1D). Importantly, this histogram contained all data from the low, medium, and high nucleosome density chromatin samples shown in Figure 1.

Autocorrelations were calculated using Python, then clustered as described below. All scripts for computing autocorrelation are available at the above link.

Leiden clustering analyses were performed identically to (*25*).

**SAMOSA-ChAAT quality control validation**

Correlation of footprint midpoints for fluorescently end-labeled *Cyp3a11* vs unlabeled *Cyp3a11* is shown in Fig. S7A. Correlation of footprint midpoints for chromatin methylated in a test tube vs methylated in a microscopy plate well shown in Fig. S7B. Correlation between SAMOSA-ChAAT technical replicates is shown in Fig. S7C.

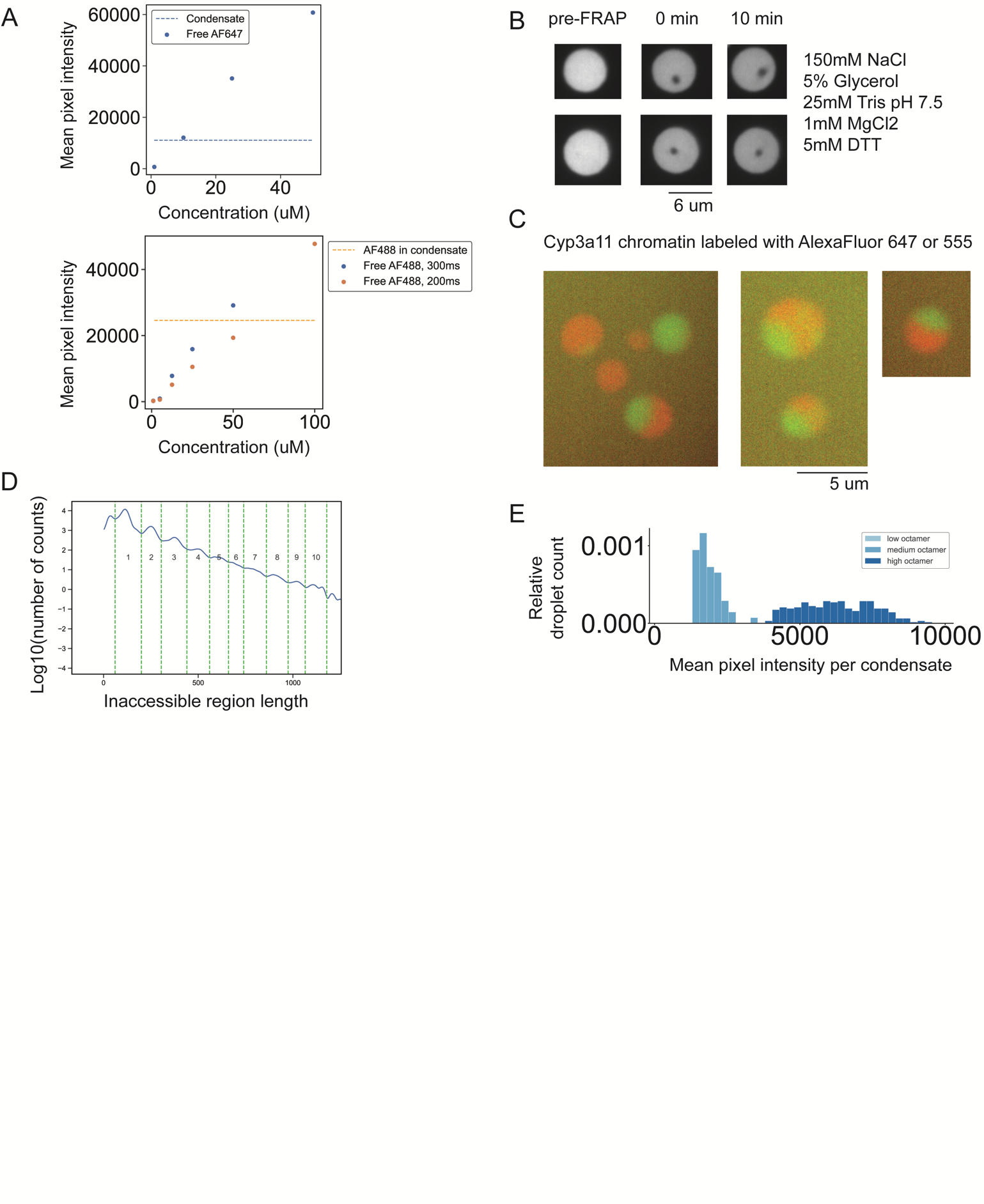

**Figure S1. Characterization of chromatin in condensates.** (**A**) Standard curves for AF647 and AF488 using the same exposure time and laser power as in experimental conditions. (**B**) Confocal images of two example AF647-Cyp3a11 chromatin condensates in the listed buffer conditions. Quantification of recovery shown in Fig S2B. (**C**) Confocal images from two-color mixing experiment using AF555 and AF647 labeled chromatin. (**D**) To estimate the number of nucleosomes on each DNA molecule, cutoffs were defined to delineate between the number of estimated nucleosomes within an inaccessible region. Green dashed lines show the cutoffs, and the numbers below indicate the number of nucleosomes that sized region is counted as. Cutoffs were based on the peaks in region length. (**E**) Histogram of mean pixel intensity per condensate for the conditions shown in Fig 1B, with histogram bin number equal to 0.1*number of condensates per condition.

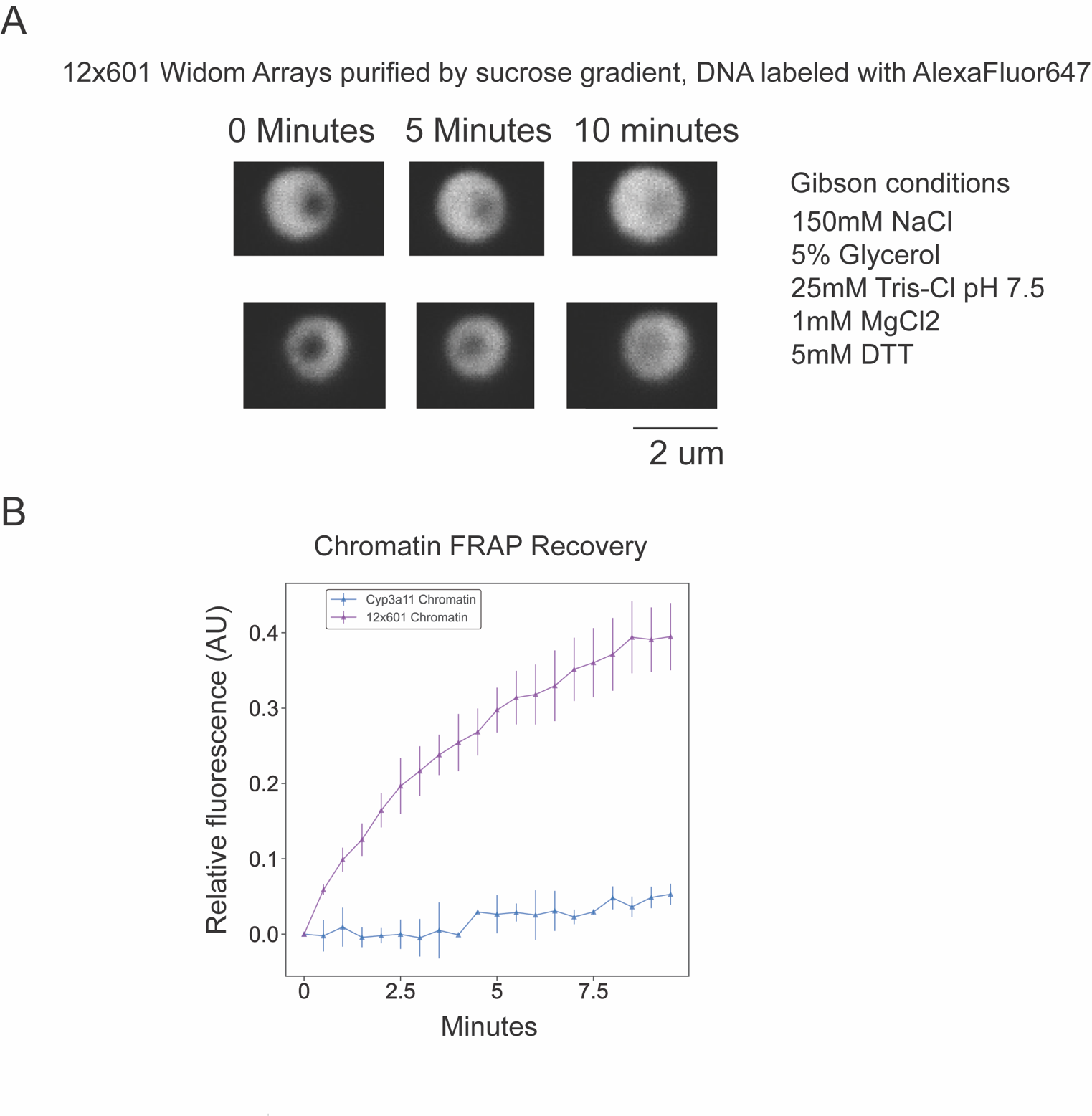

**Figure S2. Repetitive chromatin arrays recover from photobleaching.** (**A**) Confocal images of two example 12xWidom601 chromatin condensates labeled with AF647 before and after photobleaching in the listed buffer. (**B**) Quantification of recovery after photobleaching for the chromatin in A, n=5 droplets, and the chromatin in S1B, n=2 droplets. Recovery is normalized to pre-bleach droplet intensity.

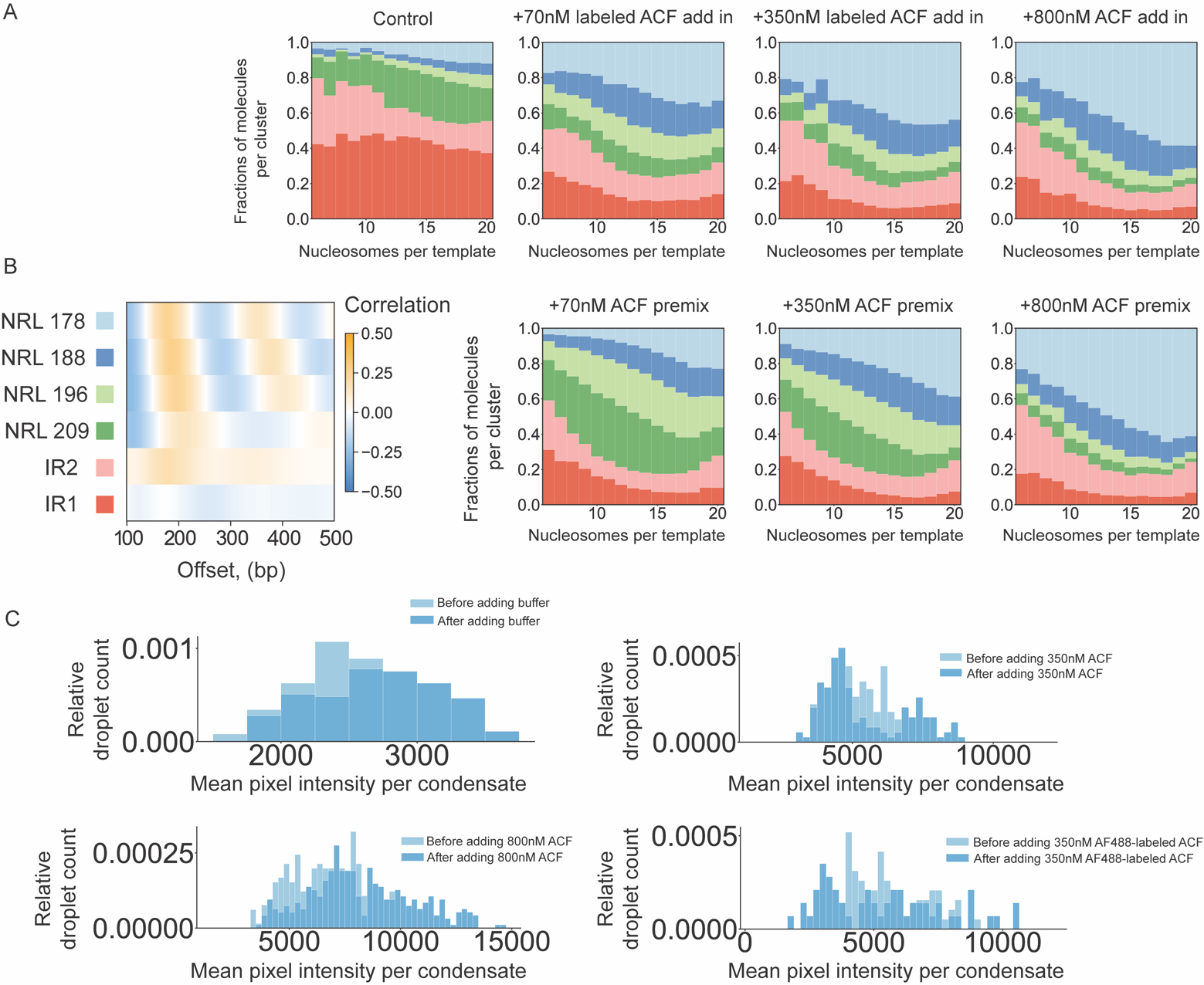

**Figure S3. ACF is a density dependent remodeler within chromatin condensates and does not reduce condensate intensity.** (**A**) Stacked bar chart representation of cluster representation in Fig 4D plotted as a function of nucleosome density. (**B**) Average single-molecule autocorrelograms following Leiden clustering of individual molecules as shown in Fig 4D. All samples were clustered together. (**C**) Histograms of mean pixel intensity per condensate before and after ACF add-ins, with histogram bin number equal to 0.1*number of condensates per condition.

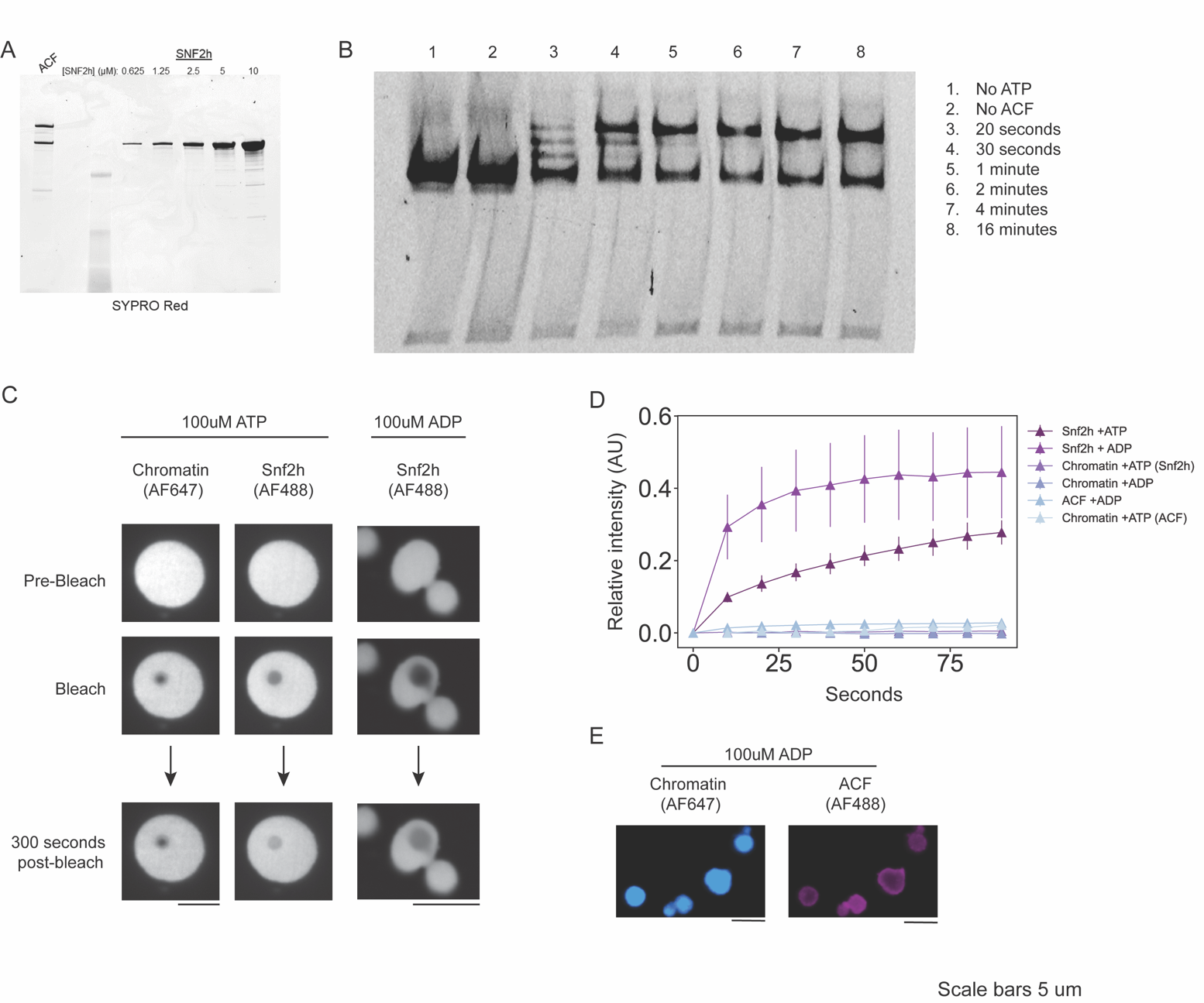

**Figure S4. Snf2h recovers from photobleaching and uniformly distributes through chromatin condensates.** (**A**) Quantification gel of ACF using Snf2h standards. (**B**) Validation that ACF centers end-positioned 0-60 Widom601 mono-nucleosomes. (**C**) Confocal images of AF647 and AF488 fluorescence in various add-in conditions. Fluorescence intensity is not normalized across columns. (**D**) Quantification of recovery after photobleaching for the condensates in C along with additional conditions, n=3-5 droplets per condition. Recovery is normalized to intensity of unbleached droplets over the time course. (**E**) ACF distribution throughout chromatin condensates in conditions with ADP.

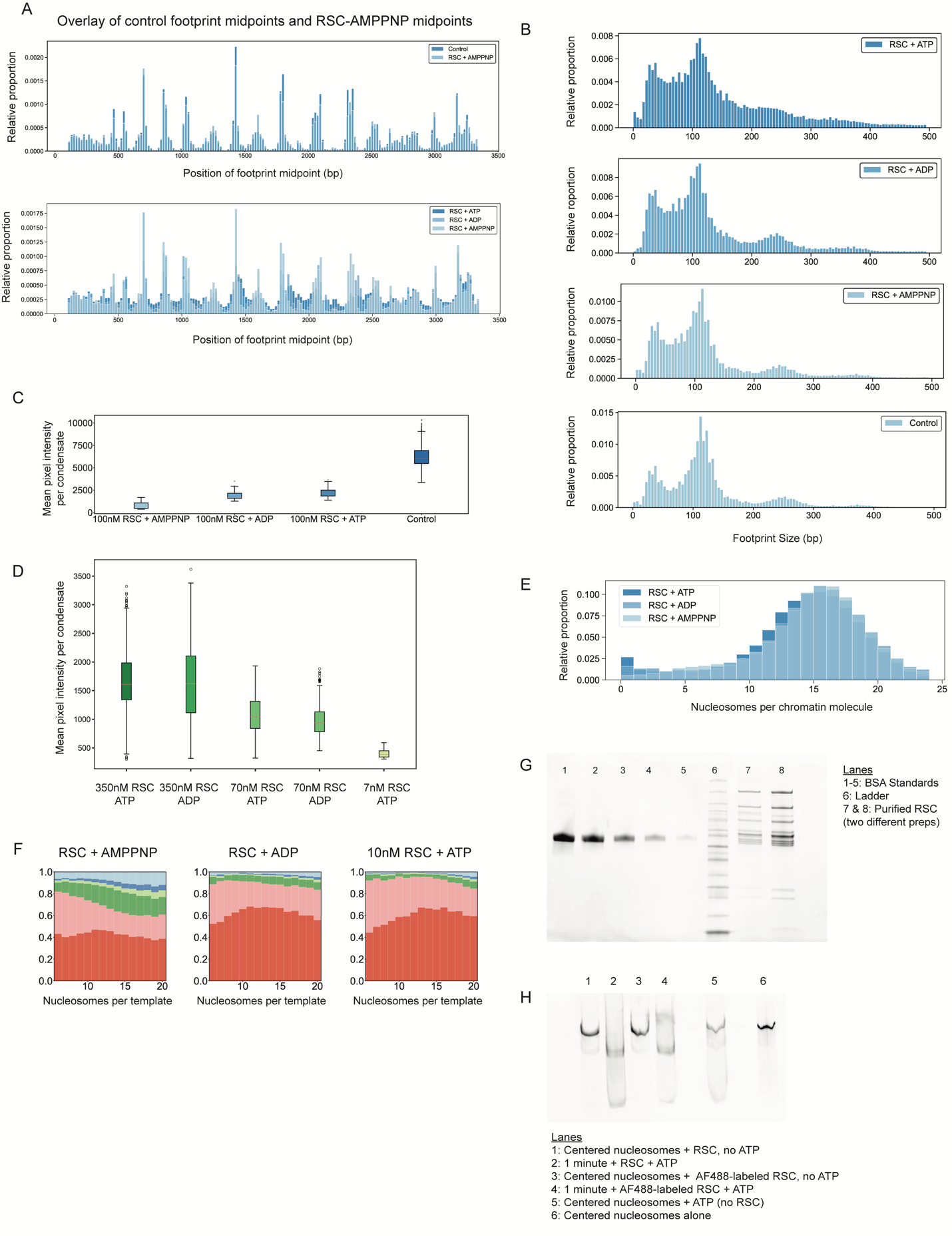

**Figure S5. Effects of RSC remodeling on chromatin.** (**A**) Histograms of footprint midpoints across the Cyp3a11 sequence, bin size 20 base pairs. (**B**) Footprint size distributions in RSC conditions depending on nucleotide. (**C**) Boxplot of mean pixel intensity per condensate (AF647 channel) in each RSC condition. (**D**) Boxplot of mean pixel intensity per condensate (AF488 channel) after adding labeled RSC into reactions. (**E**) Nucleosomes per chromatin molecule for each condition with RSC. (**F**) Stacked bar chart representation of cluster representation in Fig 4D, plotted as a function of nucleosome density. (**G**) Quantification gel of RSC using BSA standards. (**H**) Validation that labeled and unlabeled RSC remodel centered 40-40 Widom601 mono-nucleosomes.

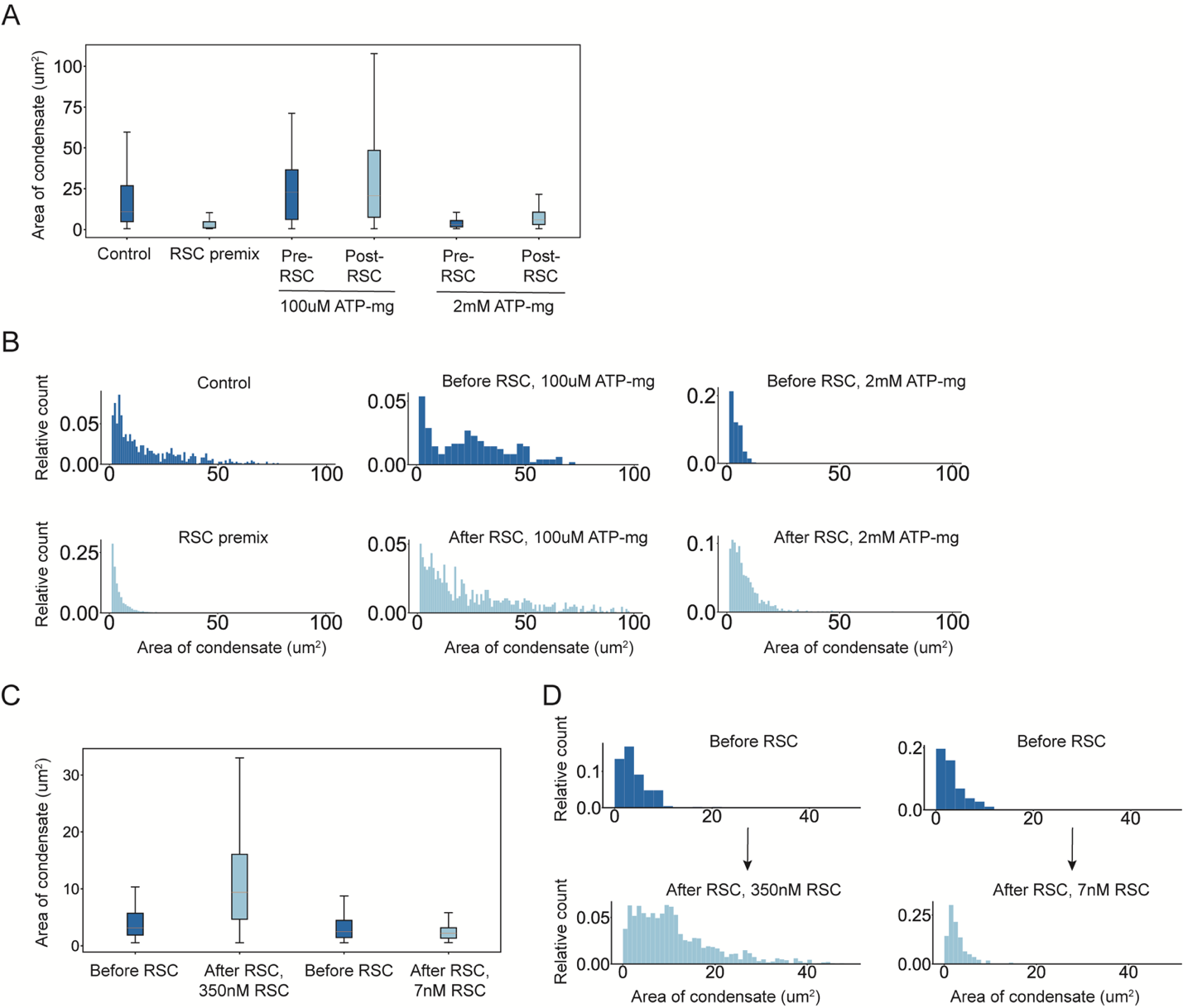

**Figure S6. Effects of RSC on condensate area.** (**A**) Boxplot of condensate area in each RSC condition. (**B**) Histogram of condensate area before and after RSC add-in, with histogram bin number equal to 0.1*number of condensates per condition. (**C**) Boxplot of condensate area in each condition with AlexaFluor-488 labeled RSC. (**D**) Histogram of condensate area before and after labeled RSC add-in, with histogram bin number equal to 0.1*number of condensates per condition.

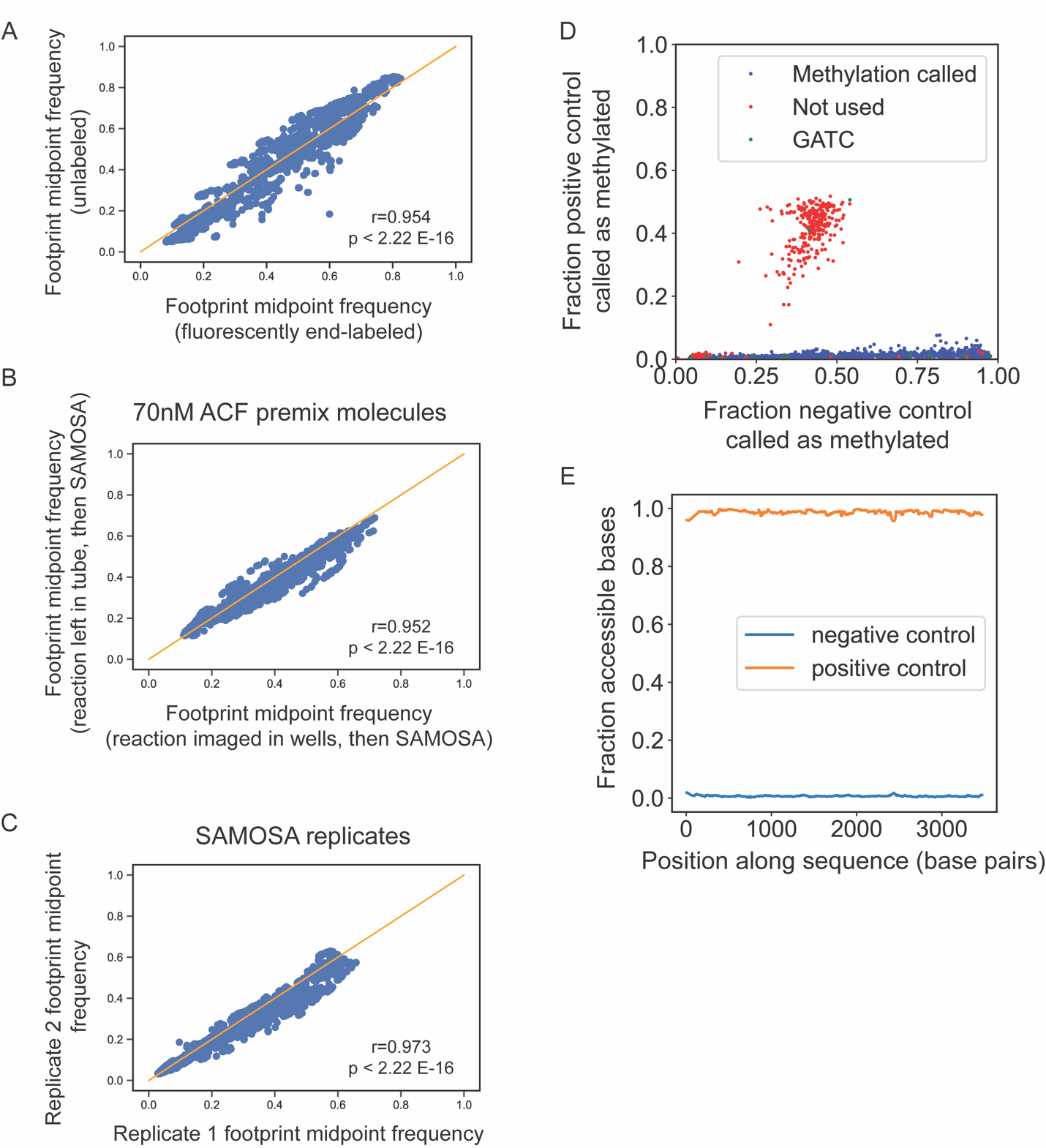

**Figure S7. SAMOSA quality control validation.** Correlation of midpoint frequency for (**A**) unlabeled chromatin and AF647-labeled chromatin, (**B**) chromatin left in tubes before SAMOSA and chromatin added to microscopy plate before SAMOSA, (**C**) two replicate SAMOSA experiments. Correlated molecules all have the same number of nucleosomes per molecule. (**D**) Adenine distribution for all adenines on Cyp3a11 sequence. (**E**) Average methylation for positive and negative control molecules (Cyp3a11 DNA fully methylated or unmethylated, respectively).

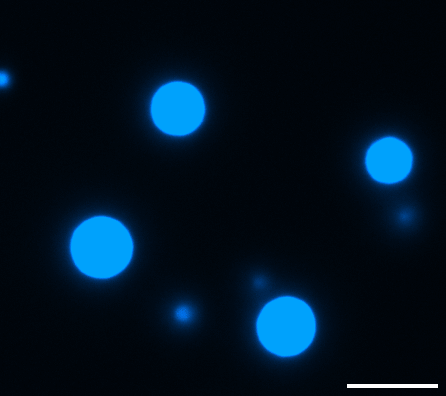

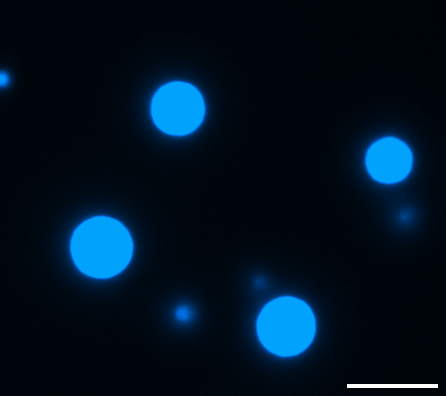

**Movie S1 and S2.** Premix control condensates, 100uM ATP-mg. Speed is one fps (left) and five fps (right). Scale bar is 10 um.

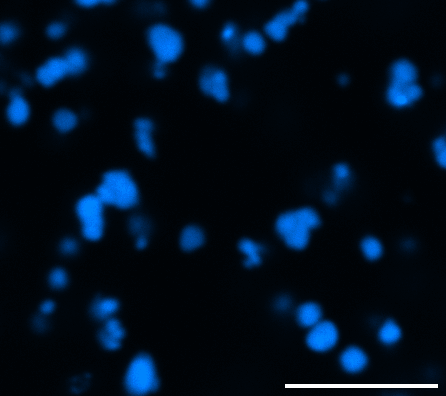

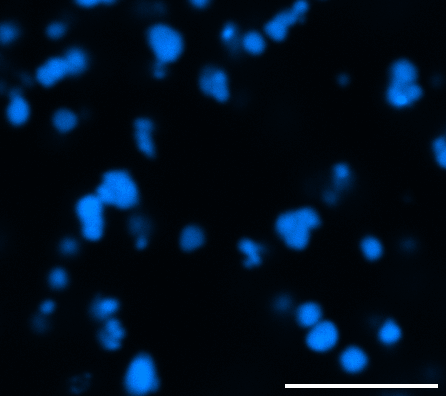

**Movie S3 and S4.** Premix condensates + RSC, 100uM ATP-mg. Speed is one fps (left) and five fps (right). Scale bar is 10 um.

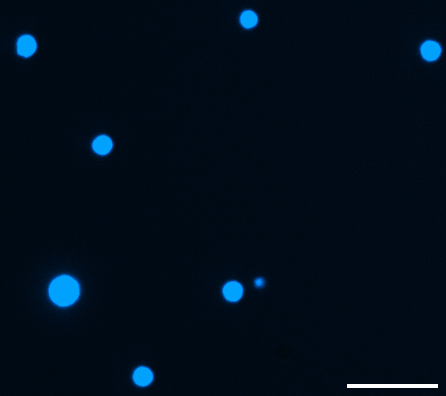

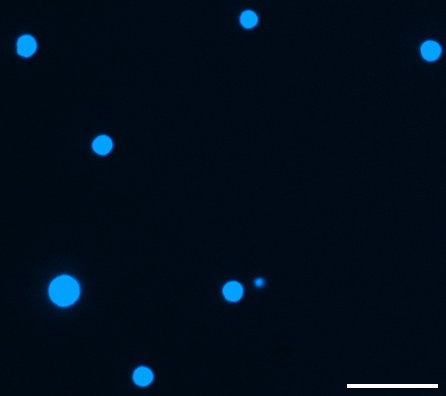

**Movie S5 and S6.** Before adding RSC to condensates, 2mM ATP-mg. Speed is one fps (left) and five fps (right). Scale bar is 10 um.

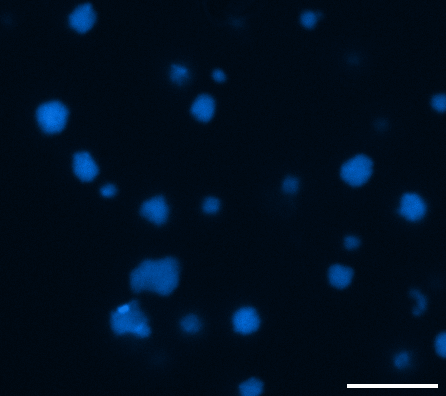

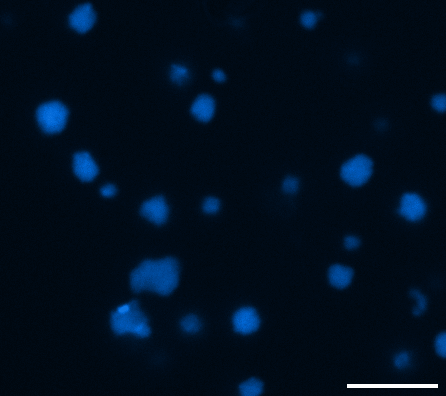

**Movie S7 and S8.** After adding RSC to condensates, 2mM ATP-mg + 500nM RSC. Speed is one fps (left) and five fps (right). Scale bar is 10 um.

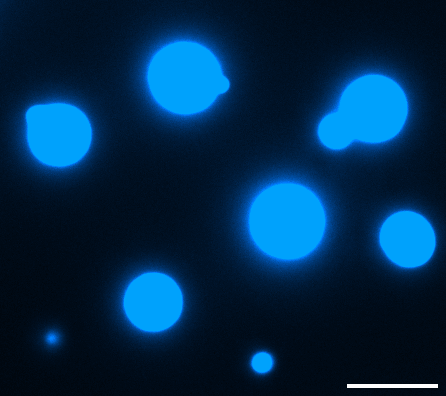

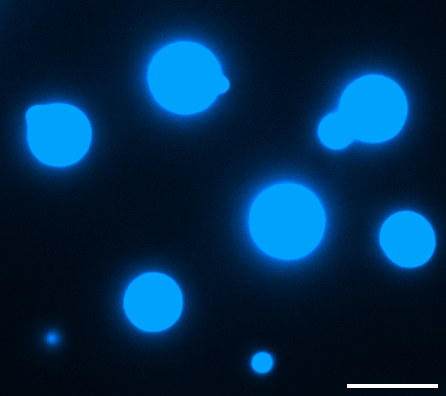

**Movie S9 and S10.** Before adding RSC to condensates, 100uM ATP-mg. Speed is one fps (left) and five fps (right). Scale bar is 10 um.

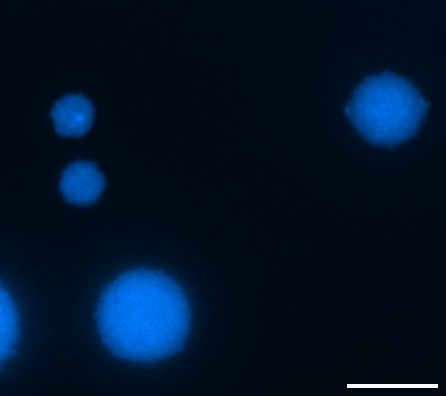

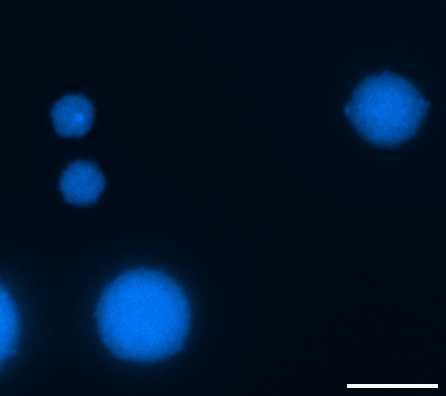

**Movie S11 and S12.** After adding RSC to condensates, 100uM ATP-mg + 500nM RSC. Speed is one fps (left) and five fps (right). Scale bar is 10 um.
